# Miniaturizing, Modifying, and Magnifying Nature’s Proteins with Raygun

**DOI:** 10.1101/2024.08.13.607858

**Authors:** Kapil Devkota, Daichi Shonai, Joey Mao, Young Su Ko, Wei Wang, Scott Soderling, Rohit Singh

## Abstract

Proteins have evolved over billions of years through extensive and coordinated substitutions, insertions and deletions (indels). Computational protein design cannot yet fully mimic nature’s ability to engineer new proteins from existing templates. Protein language models generate informative per-residue representations, but leveraging them to execute large-scale, function-preserving mutations and indels has remained beyond reach. We introduce Raygun, a generative AI framework that unlocks efficient miniaturization, modification, and augmentation of proteins, using a novel probabilistic encoding of protein sequences constructed from language model embeddings. Emulating evolution, Raygun shrinks proteins by 10-25% (sometimes over 50%) while preserving predicted structural integrity and fidelity, introduces extensive sequence diversity while preserving functional sites, and can expand proteins beyond their natural size. These capabilities unlock new opportunities in gene therapy and biotechnology. In cell-based validation, Raygun successfully miniaturized fluorescent proteins, two of which are smaller than 96% of fluorescent proteins reported in FPbase, as well as TurboID, a synthetic biotin ligase widely adopted for proteomics. It also successfully expanded Epidermal Growth Factor (EGF), a natural binding partner to the EGFR protein, generating EGF variants with higher binding affinity than the wildtype. Raygun’s conceptual innovations in template-based protein design reveal that protein function can be encoded in a length-independent space. This fundamental insight bridges protein representation learning with evolutionary biology and could unlock the development of more efficient molecular tools and biological therapeutics.

## 1 Introduction

Protein design has recently made significant strides, particularly in de novo creation of proteins tailored to specific functions or structures [42, 2, 20, 19, 23]. Yet, the landscape of protein engineering extends beyond crafting entirely new molecules. Evolution argues that a compelling alternative is to build upon existing proteins, an approach we call template-guided design (**Figure 1A-C**). This approach, akin to renovating a building rather than constructing from scratch, has wide-ranging applications such as miniaturizing proteins, modifying established reporters and sensors while maintaining their core functionality, or adapting gene payloads for viral vector delivery size constraints. Despite its potential, current template-based approaches face significant limitations. They typically rely on point substitutions—essentially swapping out individual amino acids—and struggle to incorporate more extensive modifications. As substitutions increase, the combinatorial space explodes exponentially (e.g., 25 locations yield 10^32^ possibilities), making computational prediction and experimental validation impractical. Beyond combinatorial substitutions, current methods are fundamentally unable to fully mimic nature’s mutational repertoire. Natural evolution generates new proteins not just through substitutions, but also via insertions and deletions (indels). Modifying proteins without the ability to incorporate indels is akin to renovating a building without the ability to add or remove entire rooms. Unfortunately, the field lacks principled, scalable methods capable of accommodating substantial indels while maintaining a protein’s core structural form. The potential impact of overcoming this hurdle is profound: a design approach that could manage both combinatorial substitutions and large-scale indels would vastly expand the universe of proteins derivable from a single template.

**Figure 1:**
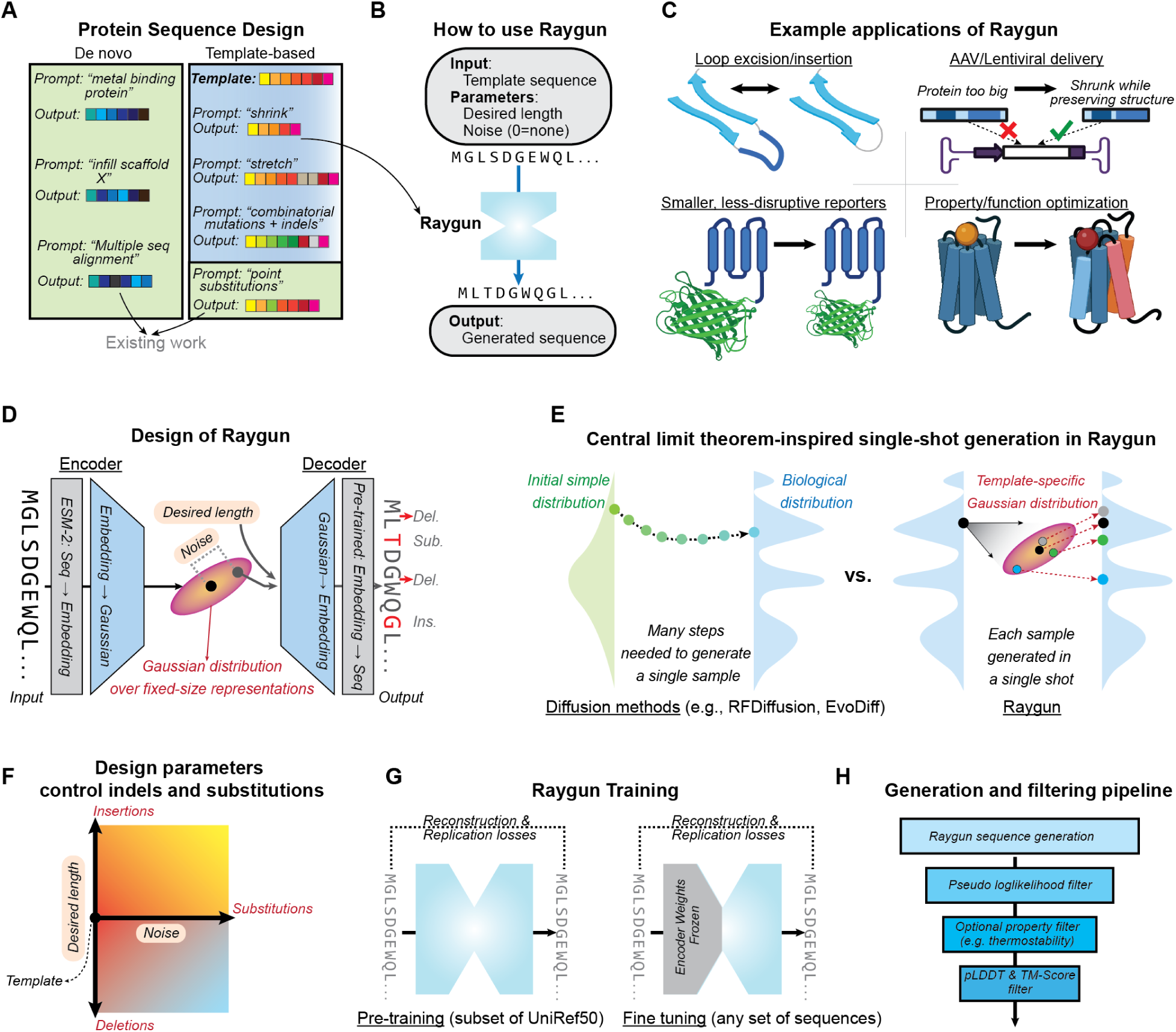
Description of the Raygun Model. **(A)** Unlike de novo models, which design entirely novel proteins conditioned on structural or functional constraints, Raygun uses existing proteins as a template and introduces appropriate substitutions and indels in a template-guided design. **(B)** It takes only three inputs: the template sequence, a noise parameter and target length. **(C)** Raygun has direct and wide-ranging applications in biological research, ranging from protein sensor design to protein miniaturization for gene therapy. **(D, E)** It works as an auto-encoder model. The encoder converts variable-length sequence input into a fixed-dimensional representation, and this space approximates a multi-variate normal distribution. Unlike diffusion-based models, single-shot sampling can be done through this distribution. **(F)** Raygun parameters provide fine-grained control over indels and substitutions. **(G)** It is trained in a self-supervised way. For inference, we recommend using a fine-tuned version in which only the decoder block is updated. **(H)** Raygun is fast (0.3 seconds per generation) and can be paired with downstream filtering steps to generate high quality candidates efficiently.

Protein language models (PLMs) offer a promising avenue for addressing these challenges. By representing proteins as sentences of amino acids, PLMs harness evolutionary conservation to learn distributional rules, generating rich, high-dimensional embeddings of protein sequences: a sequence of *n* amino acids maps to *n* points in a high-dimensional space, with each point encapsulating both local and global context of its corresponding residue [24, 27, 21, 6, 14]. PLM embeddings have proven remarkably powerful: they have been applied to predict protein interactions, structures, and functional properties [37, 34, 35, 21, 24, 44]. From a protein design standpoint, the PLM embedding space presents an enticing blend of representational fidelity and computational efficiency. For point substitutions, sampling in the neighborhood of each residue’s embedding representation has been shown to generate residue substitutions that preserve peptide binding [4]. We hypothesize that PLMs can be leveraged for template-guided design by reducing the difficult task of sequence-space design to the easier task of embedding-space design. Importantly, sequence–embedding conversions are highly accurate. We use ESM-2, a PLM pre-trained on millions of sequences, for sequence-to-embedding transformation. For the reverse transformation, we trained a small neural network that achieves over 99% validation accuracy; others report similarly high performance [12, 8]. These bidirectional transformations allow us to operate in the structurally-aware embedding space while maintaining a clear path back to realizable sequences, opening new design possibilities. However, the critical challenge of handling indels and combinatorial substitutions remains to be addressed: PLM embedding size varies with sequence length, making proteins of different sizes incompatibly represented in different-dimensional spaces.

Our key conceptual advance is to encode each protein not as a point in high-dimensional space, but as a probability distribution. To construct this probabilistic encoding, we introduce a fixed-size PLM representation addressing two critical challenges: a) efficient, principled sample generation, and b) bidirectional translation between this representation and variable-length embeddings (and hence sequences). Specifically, we represent any protein of length 50 or higher as a 64,000-dimensional multivariate normal (MVN) distribution. Our design strategy generates candidates by directly sampling from the template protein’s MVN distribution. This approach enables “single-shot” design: unlike diffusion-based methods [38, 39, 42, 22] where output quality and diversity depend on intermediate denoising steps, our method generates high-deviation candidates just as efficiently as low-deviation ones, and much faster than diffusion-based methods. We drew inspiration from the central limit theorem (CLT): The CLT states that the sum of identically distributed random variables, given appropriate independence criteria, approaches a normal distribution [13, 32]. We partition proteins of length 50 or greater into a fixed number of segments, each characterized by its own MVN distribution. These segment-level distributions collectively form the full protein’s distribution.

We introduce Raygun, a deep learning framework implementing these concepts. An encoder-decoder architecture, Raygun unlocks the generation of diverse, high quality candidate sequences from a given template (**Figure 1**). It offers broad flexibility to the protein designer, allowing them to specify the extent of shrinkage, stretching, and substitutions by two intuitive parameters: the desired output length and the level of noise. The noise parameter broadly controls the substitution rate; indels can be calibrated by changing the output length.

Raygun generates candidates with the desired level of deviation– small or large– from the template while preserving predicted foldability and structure. We observed that sequences of most proteins could be stretched or shrunk by as much as 10% with modest structural impact (median pLDDT [43] decrease against template ∼ 17%, TM-score [45] ∼ 0.78). At the same time, Raygun can also generate large deviations, such as halving or doubling the protein length or substituting over 50% of the residues, all while broadly preserving the original computed structure. Raygun’s speed is valuable during generation: its single-shot generation is 100-fold faster than diffusion-based de novo design approaches and pairing it with PLM-based evolutionary likelihood filters enables us to generate candidates that have large deviations from the template but nonetheless with attractive foldability and structural properties. Across diverse PFAM [3] families, Raygun consistently produces high-quality sequences, preserving PFAM motifs (48.5%) even when substantially altering protein length (50-200% of original length). Unlike some de novo approaches, Raygun’s performance is consistent across different secondary structure elements, handling both beta sheets and alpha helices effectively.

As demonstration, we applied Raygun to generate novel fluorescent proteins variants [33, 18] with properties not available in any protein currently characterized, natural or artificial. Fluorescent proteins (FPs) are essential tools in cell biology, but their size may disrupt the functions of proteins they’re fused to, especially small proteins. Using Raygun, we generated 8 miniaturized candidates across two FP templates. Five of these candidates showed fluorescent activity matching the spectrum of the original template, albeit at lower intensity. Notably, two of the generated proteins, at lengths of 199 and 206 amino acids, both derived from mCherry, are shorter than 96% of fluorescent proteins reported in FPbase [18]. One of these deviated from the chromophore sequence of its template (mCherry), underscoring Raygun’s potential to synthesize new-to-nature designs.

We further evaluated Raygun’s ability to edit multi-domain proteins with enzymatic activity by generating miniaturized variants of TurboID, a synthetic biotin ligase derived from BirA. Experimental validation showed that 55% (6 out of 11) of the designed variants were successfully expressed in cells, with two exhibiting ligase activity. This highlights Raygun’s proficiency in template-guided modifications, even for complex multi-domain sequences. Finally, to assess Raygun’s ability to magnify proteins, we used it to generate enlarged variants of the wildtype Epidermal Growth Factor (EGF) protein and measured their binding affinity to the Epidermal Growth Factor Receptor (EGFR). All four tested candidates were successfully expressed, with two displaying higher binding affinity than wildtype EGF. While Raygun is not specialized for binder design, its ability to enhance binding affinity without task-specific modifications underscores its potential as a general-purpose tool in protein engineering and therapeutic design.

Raygun’s high hit-rate, both computationally and experimentally, combined with its speed and tunability, unlocks computational avenues in protein design and engineering. Conceptually, Raygun represents a novel application of protein language models, utilizing a rich fixed-size representation that extends beyond discriminative learning to enable generative tasks. This approach has the potential to accelerate protein engineering across a wide range of applications, from developing more efficient molecular tools to designing novel therapeutics.

## 2 Results

### 2.1 A fixed-sized language for all proteins

Constructing a model that accepts variable-length PLM embeddings and generates viable candidate sequences of any length presents significant design challenges. Our objective of allowing the user any choice of output sequence length necessitates a fully length-agnostic protein representation at some internal stage. The generative model must robustly sample in this length-agnostic space, and subsequent stages must map the fixed representation back to variable lengths accurately. To address these constraints, we designed Raygun as an autoencoder. The encoder performs the variable-to-fixed transformation, while the decoder reverts the fixed-length embedding to the variable length space.

Ensuring effective variable-to-fixed mapping in the Raygun encoder was crucial to the model’s success. ESM-2 embeddings, trained via masked language modeling (MLM), provide local and global protein information at each residue. We hypothesized that these variable-length embeddings have enough redundancy to allow fixed-length compression with minimal information loss. To address this, we introduced a two-level representation. First, we modeled any variable-length ESM-2 embedding as a series of *K* contiguous blocks, focusing on template proteins of at least 50 residues. We chose *K* = 50 after a preliminary comparison between *K* = 50 and *K* = 25 (**Figure A.3**). We posited that the series of *K* blocks could be trained to recapitulate the template’s global representation. Each block’s size depends on the template protein length (e.g., 10 for a 500-length template), but we envisioned that the per-residue embeddings have enough redundancy to be reduced to a fixed-size representation without significant information loss. By the Central Limit Theorem for weakly dependent variables [32], averaging across a block’s length should approximate a Multivariate Normal Distribution (MVN) which can be sampled directly. Thus, the Raygun encoder can produce a fixed-length embedding space that is tractable for sampling and retains sufficient information for the Raygun decoder to accurately map it back to variable lengths.

### 2.2 Raygun architecture & training

The Raygun architecture consists of two length-transforming layers, “Reduction” and “Repetition,” and multiple length-preserving “T-Block” layers. The Reduction and Repetition layers are parameter-free and operate in the encoder and decoder stages, respectively. T-Block layers, present in both stages, are deep neural network modules trained to optimize embeddings by enhancing latent local and global properties. Each T-Block layer comprises an ESM Transformer (hence the ‘T’) for global properties and a 1D-convolution block for local relationships. They are used before, between, and after the length-transforming layers (**Methods A.1**). T-Block parameters comprise most of Raygun’s 701 million trainable parameters.

In each Reduction layer, within-block averaging generates two fixed-length outputs per block: the block-wide mean and standard deviation matrices, describing the MVN distribution of the fixed-length space. During inference, the user-supplied noise scale dictates the sample generated from this MVN for subsequent layers; during training, the MVN’s mean is used. The Repetition layer accepts a target length and a fixed-length representation from the encoder stage, producing a variable-length embedding of the desired length.

#### Training the Raygun model

We formulated Raygun training as a self-supervised problem. Specifically, the model is trained to compress the input sequence to a fixed size, then decompress it back to its original length with maximal fidelity. We assessed the quality of the reconstructed output in both the embedding and sequence spaces, by using two losses: (a) reconstruction loss, penalizing deviations in the reconstructed embeddings, and (b) cross-entropy loss, penalizing deviations in the sequence space after using a pre-trained ESM-2-decoder. Additionally, we sought self-consistency in the fixed-length embedding. For each input sequence, we decoded the fixed-length representation to a shorter-length sequence, re-encoded it, and compared the two fixed-length representations under a replication loss (**Methods A.2**). Together, these three losses comprised the overall training objective.

We paid special attention to curating the training data, reasoning that good generalizability would benefit from a wide, even distribution of protein lengths. After inspecting the length distribution in Uniref-50, we focused on proteins of lengths between 100 and 1000, dividing this range into 19 bins. For each bin, we randomly selected roughly 5000 sequences within the specified length range from the Uniref-50 database, resulting in a total of 94,734 proteins. Finally, of these extracted Uniref-50 proteins, we randomly chose 80,000 proteins for model training and used the remaining 14,734 proteins for validation and hyper-parameter selection. During training, we computed the BLOSUM-weighted [10] sequence identity score (“BLOSUM score”) at each epoch, and chose the epoch with the highest validation score as the pre-trained model for fine-tuning and further experiments.

An alternative to this self-supervised approach would be supervised training where pairs of sequences with differing lengths but identical structural and functional properties are used. Unfortunately, this is an infeasible approach. We could not find experimental datasets where large-scale indels were performed with this goal in mind. Databases like SCOP [25] and CATH [26], while grouping proteins by structural domains, are domain-focused rather than ensuring structural and functional similarity at the whole-protein level.

#### Raygun fine-tuning and sampling

Akin to pre-trained foundation models that are fine-tuned for specific use-cases, we recommend fine-tuning Raygun before a protein design campaign. This can be done simply and efficiently. For a single protein, or a group of proteins (e.g., the relevant PFAM domains), a simple FASTA file can be provided as input to fine-tune the autoencoder; no alignment is needed, and diverse sequences can be included. The fine-tuning is designed to improve the fidelity of Raygun decoder for the particular protein group, so that the decoder can accurately transform the fixed-length representation back to the variable length sequence space with BLOSUM score *>* 0.99. To avoid the risk of catastrophic forgetting seen sometimes in fine-tuning and noting that the actual sampling is controlled by the encoder segment of Raygun, we froze the encoder weights while doing fine-tuning optimization; only the decoder is fine-tuned to ensure the generation is well-matched to the proteins of interest.

Although Raygun can perform single-shot protein generation (unlike other diffusion-based models), we found that using a one-step recycling iteration improves performance. Like the diffusion process, where an intermediate output is again passed to the input, recycling in Raygun takes the generated candidate and uses it as a template to generate a new candidate. We noted that it improves the quality and diversity of output samples (**Method A.3.5**).

Our generation process is fast enough (0.3 secs/iteration on an A100 GPU) that, even after onestep recycling, we can generate a large number of viable candidates for moderately sized proteins of length *<* 1000. Separately, we note that Raygun works well even on very large proteins (like mTOR), albeit slower. Raygun’s speed gives us the flexibility to apply filtering schemes to select higher-quality candidates from the generated samples. We primarily used a PLM-based metric called pseudo loglikelihood (*pLL*), which assesses the fitness of a protein as modeled by language models like ESM-2 [5]; we devised a length-adjusted version (**Method A.3.1**). For each generated sample, we employed *pLL* to rank the candidates by their scores, selecting those with the highest fitness. While other PLM and structure-based filtering techniques were used in some experiments, *pLL* was consistently the first filtering step due to its computational efficiency.

### 2.3 Raygun preserves protein structure while introducing indels & combinatorial mutations

Raygun effectively generates novel candidates across a wide range of template sizes while largely maintaining the candidates’ structure and integrity. As a demonstration of this capability, we applied Raygun to a diverse set of proteins (**Figure 2**), including hemoglobin (147 aa), CCR1 (355 aa), lacZ (or BGAL ECOLX; 1029 aa), and mTOR (2549 aa). Our goal was to generate both shortened and elongated variants of these proteins, assessing Raygun’s ability to introduce insertions, deletions, and substitutions while preserving overall computed structure.

**Figure 2:**
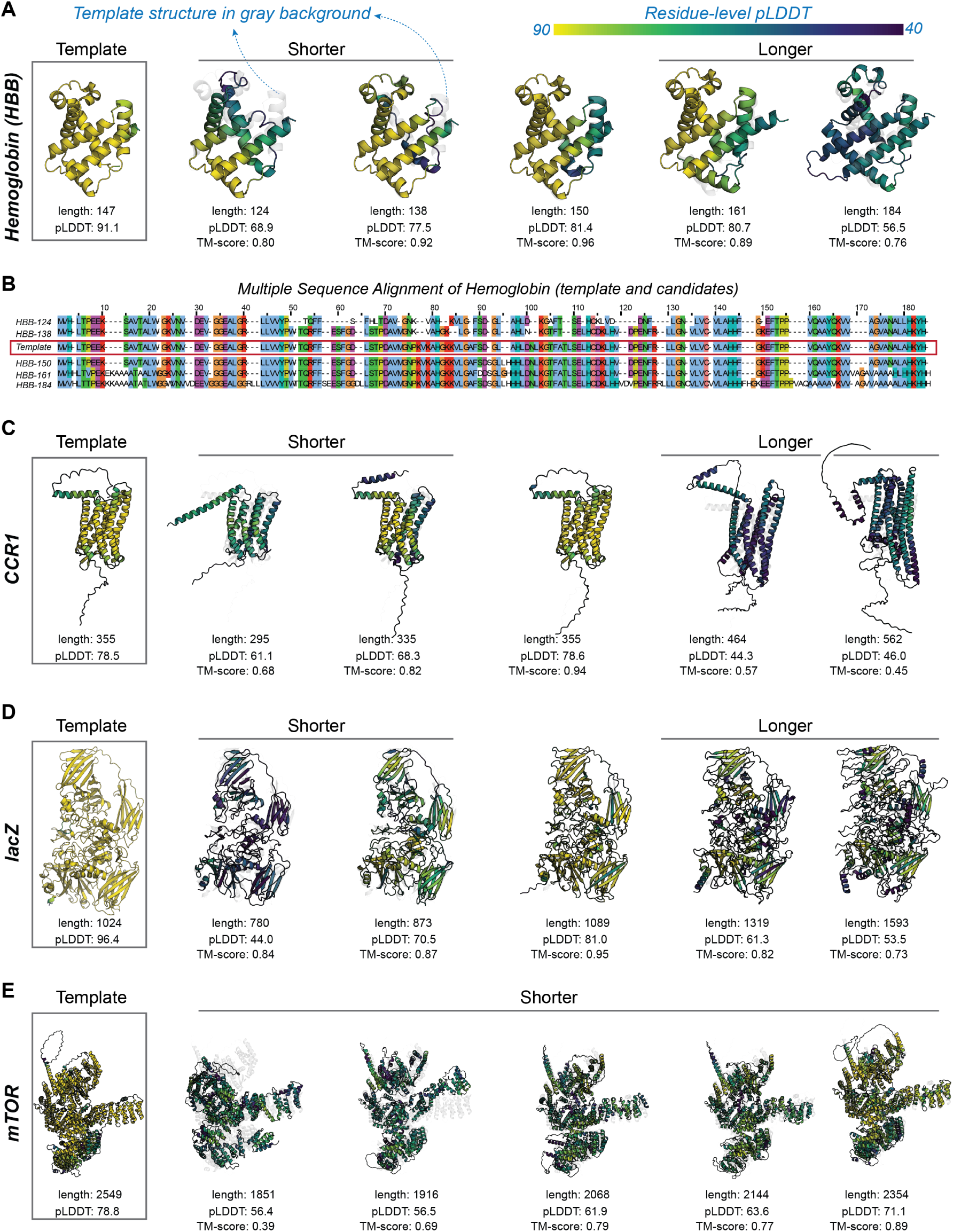
Protein editing using Raygun for proteins of different sequence lengths. The following proteins were chosen for a demonstration of Raygun’s miniaturization & magnification capabilities: **(A)** Hemoglobin (HBB), **(C)** CCR1, **(D)** lacZ and **(E)** mTOR. For HBB, the multiple sequence alignment of the Raygun candidates and the templates are shown in **(B)**. For HBB, CCR1 and lacZ, we showed both the shrunk and enlarged results. For mTOR, however, owing to its large size, we show only its selected shrunk candidates.

We generated 2000 samples for each protein across a broad range of lengths, setting Raygun’s sampling noise parameter to 0.5 to introduce moderate sequence variations. To ensure the quality of our candidates, we filtered the samples using length-adjusted ESM-2 pseudo-loglikelihood (*pLL*) scores. We retained the top 5% of generated sequences distributed across the length spectrum (**Methods A.3.1**).

Figure 2 shows the AlphaFold-3 predicted structures of five representative candidates for each protein, along with their predicted Local Distance Difference Test (pLDDT) scores and alignment with the template (TM-scores). The results demonstrate that Raygun effectively preserves protein structure across a wide range of output lengths, as evidenced by the high TM-scores when comparing generated structures to their respective templates.

Raygun’s performance extends from small to large and complex proteins like mTOR (2549 aa). However, we observed that the degree of structure preservation varies depending on the properties of the template protein. For instance, among the samples with inferred AlphaFold-3 [1] structures, CCR1’s TM-score decreased to 0.68 when shortened by 17%, while mTOR could accommodate a 25% reduction to reach a similar TM-score of 0.69. As expected, greater deviations from the original length generally correspond to lower TM-score and pLDDT values. This trend reflects the challenge of maintaining structural integrity as proteins undergo more substantial modifications. While we used a moderate noise-factor of 0.5 for these samples, we will shortly show how variations in this parameter control sequence and predicted structural similarities of the generated proteins.

#### Framework for evaluating template-guided design methods

To establish a comprehensive framework for evaluating template-guided protein design methods, we propose a set of critical capabilities that such tools should possess. These criteria not only guided our assessment of Raygun but we expect will also be useful for future template-based design approaches.

- *Structural versatility* : An ideal method will generate diverse sequences without bias towards specific secondary structures. Recent protein design methods show a concerning bias towards *α*-helical designs [2].
- *Functional site preservation*: Maintaining critical functional regions is important in protein engineering. We assessed Raygun’s capacity to retain known active and binding sites of template proteins in the generated samples, ensuring that key functional properties are preserved.
- *Tunable modification range and sequence diversity* : A versatile design tool should enable both minor tweaks and major alterations to the template, introducing indels and substitutions to produce a spectrum of variants with high sequence diversity while maintaining essential structural characteristics.
- *Scalability across protein sizes*: An effective method should perform consistently across a range of protein sizes. This criterion ensures broad applicability in diverse protein engineering scenarios.
- *Information-rich representation*: The quality of the fixed-length representation used in the design process is crucial. It should encapsulate sufficient structural information to guide accurate sampling. We evaluated this by measuring the representation’s ability to cluster structurally related sequences and compared it to baseline ESM-2 embeddings.

In assessing Raygun’s performance on these criteria, we have sometimes focused our analyses below on Raygun-generated candidates that were shortened relative to their templates. This approach allowed for straightforward sequence alignment-based evaluation metrics. We posit that these assessments will generalize to expanded candidates, given that Raygun employs the same underlying mechanism for both shrinking and expanding proteins.

#### Raygun miniaturizes proteins with a slight preference for deleting less-structured regions

To evaluate Raygun’s structural versatility, we analyzed its behavior when miniaturizing proteins across various secondary structure elements. From the PDB database, we selected 10,000 template proteins belonging to SCOP [25] families representing *α*, *β*, *α* + *β*, and *α/β* structural classes. For each template, we generated shortened variants using Raygun and performed pairwise alignments between the miniaturized candidates and their original templates.

We categorized secondary structure elements (SSEs) into three groups: *α* helices, *β* sheets, and loops. Here, “loops” encompass all regions not annotated in the PDB file as *α* helices or *β* sheets, including intrinsically disordered regions. By examining the gaps in these alignments, we could infer which structural elements Raygun preferentially removes during the miniaturization process (**Figure 3A**).

**Figure 3:**
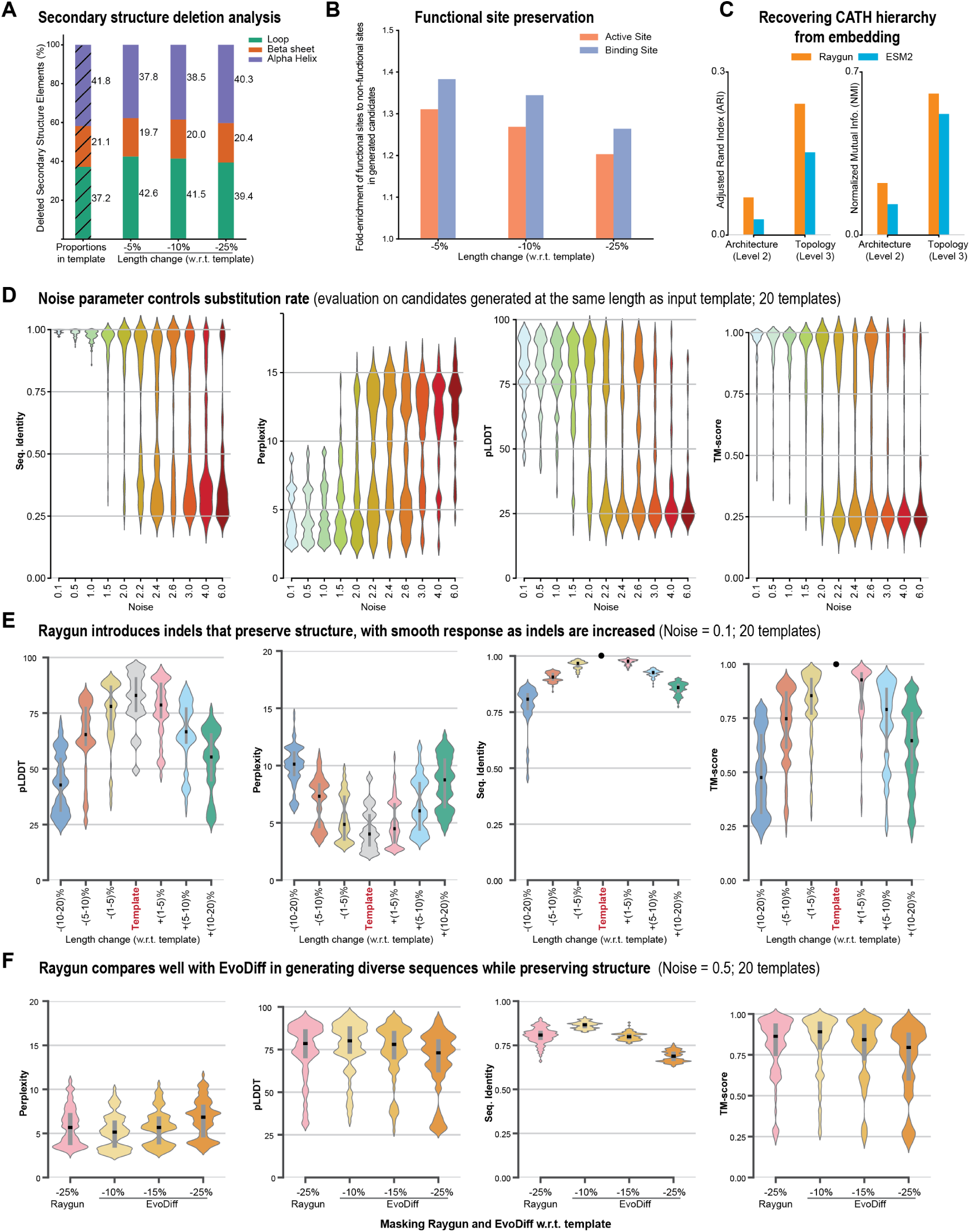
Evaluation of Raygun’s template-guided design: **(A)** Deletion preference of Raygun for secondary structure elements (SSEs). Here, “loops” refers to all non α or β residues. Raygun operates equitably across SSEs, with a slight preference for deleting loops. (*w.r.t.* = with respect to). **(B)** Ratio of Raygun preserved functional sites to the overall sequence preservation, during deletion. The ratio is consistently greater than 1, indicating that Raygun preferentially preserves active and binding sites **(C)** Clustering comparison between ESM-2 and Raygun’s fixed-length embeddings. The ARI and NMI scores show that the fixed-length embeddings produce clusters that are better aligned with the CATH structural hierarchy. **(D, E)** Structural and sequence measures for Raygun candidates at different noise and length-modification regimes. (D) The noise parameter controls substitution rates. (E) Proteins can be stretched/shrunk by upto 10% with modest loss in predicted structural fidelity. **(F)** Same-length comparisons with EvoDiff, where a part of the sequence has been masked and the methods are tasked with regenerating them. At the same masking rate as EvoDiff, Raygun’s candidates have better predicted structural properties.

Our analysis reveals that Raygun is broadly equitable in removing various SSEs, with a slight preference for removing less-structured regions. For candidates shortened by 5%, 42.6% of the gaps in the pairwise alignment corresponded to loops, although loops comprised only 37.2% of all secondary structures in the original templates. This indicates a modest bias of approximately 5% towards removing loop regions. As the degree of miniaturization increases, this preference becomes muted, leading to an even greater balance in deletions. At 25% length reduction, loops account for 39.4% of removed regions, only a 2.2% increase over the original templates. The relative propensity to remove *α* helices vs. *β* sheets remains broadly unchanged.

These results demonstrate Raygun’s structural versatility, showing that it can effectively miniaturize proteins while maintaining a balanced approach to different secondary structure elements. Its slight preference for deleting loops aligns with the general understanding that such regions often contain fewer structurally critical elements compared to *α* helices and *β* sheets. Since functional roles can be distributed across different structural elements, we next examined how well Raygun preserves known functional sites during miniaturization.

#### Raygun preferentially preserves active and binding sites during miniaturization

We analyzed 10,000 proteins from UniProt with annotated active and binding sites to assess Raygun’s ability to preserve functionally important regions during miniaturization. For each protein, we generated candidates at 5%, 10%, and 25% length reductions and performed pairwise alignments with their templates. We compared the preservation of active and binding sites to overall sequence conservation. Specifically, we calculated the ratio of preserved functional sites to the overall sequence identity between the miniaturized candidate and its template. A ratio greater than 1 indicates that functional sites are conserved more than would be expected by chance. Figure 3B shows that this ratio consistently exceeds 1 across all deletion settings, with the highest preservation observed at 5% length reduction. Binding sites are retained slightly more than active sites. These results demonstrate that Raygun effectively captures and prioritizes functionally important regions during the miniaturization process, even without explicit annotation of these sites in its training data. As expected, preservation decreases with more extensive deletions, underscoring the challenge of maintaining functional integrity during major structural modifications.

#### Raygun parameters provide fine-grained control over protein generation

Raygun offers precise control over protein generation through two key parameters: noise and output length. The noise parameter scales the covariance matrix of the multivariate normal distribution from which we sample, allowing fine-tuned control over sequence variability. To evaluate the impact of these parameters, we conducted experiments on 20 proteins spanning different structural classes (these are the same template proteins as used in the PFAM analysis in Figure 4).

**Figure 4:**
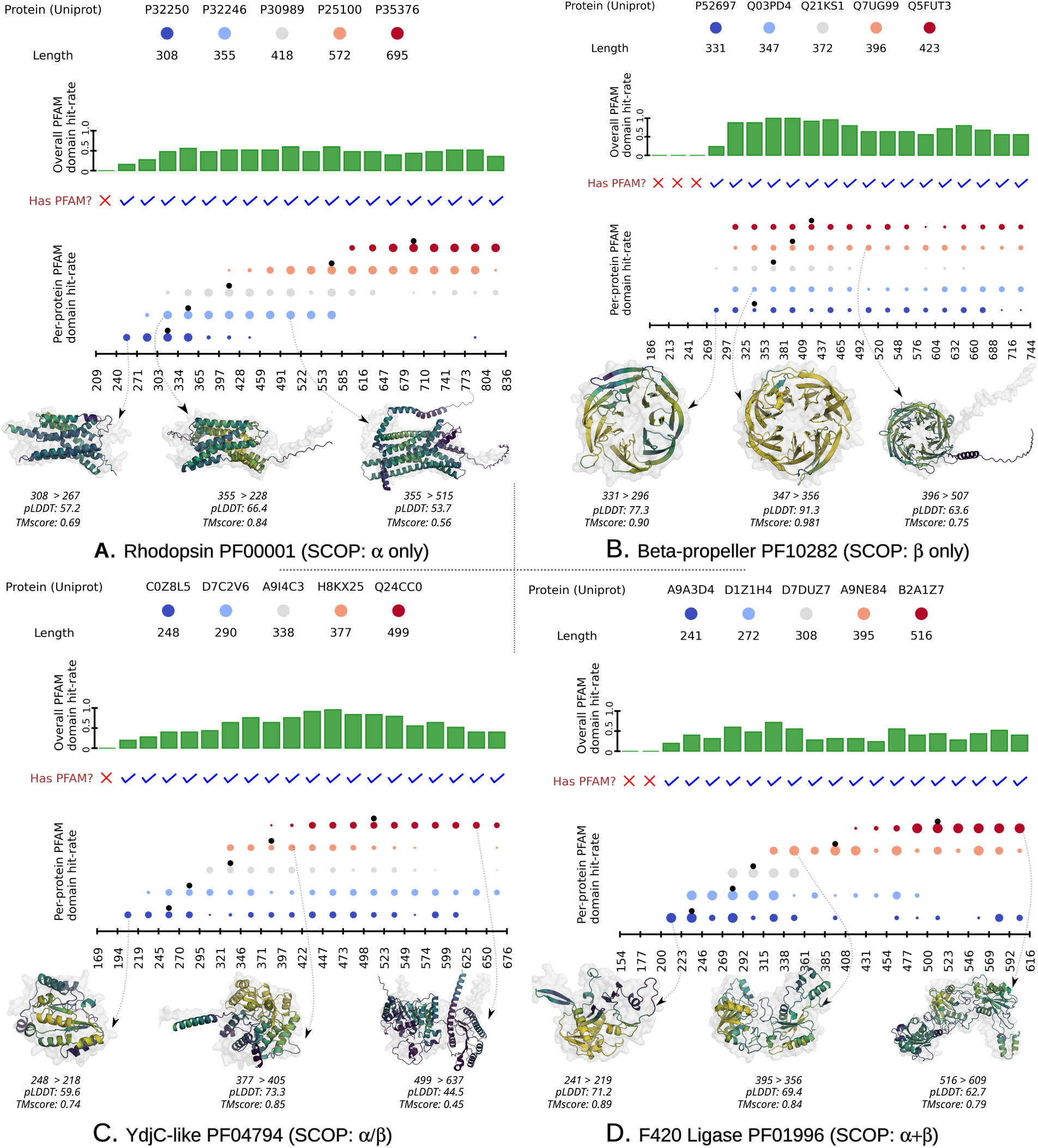
Demonstrating Raygun’s effectiveness in preserving diverse structural features, as indicated by PFAM domains. We evaluated across all four major SCOP classes: **(A)** α (*PF00001* ), **(B)** β (*PF10282* ), **(C)** α/β (*PF04794* ), **(D)** α + β (*PF01996* ). For each PFAM domain, we obtained 5 representative proteins of diverse sizes and used Raygun to generate 100 samples (per template) spanning a wide length interval (50-200% of median family length). After dividing the overall interval into 20 uniform length-bins, we report the number of Raygun candidates with retained PFAM domains across each bin. Additionally, for a selected number of candidates in each domain, we also provide their AlphaFold-3 inferred structures and metrics (pLDDT and TM-score). The template protein’s structure is shown as gray background.

First, we assessed the effect of noise on sequence and structural properties while maintaining the template length (**Figure 3D**). For each template, we generated samples for noise values ranging from 0.01 to 6, producing 100 candidates per setting and retaining the top 5 after filtering. As noise increases, sequence identity gradually decreases until an inflection point around 2.2, after which the decline becomes steeper. Structural metrics like TM-score (against template) and pLDDT follow similar trends. These results suggest using noise values near 0 for minor edits and approaching 2 for greater sequence variability. Next, we examined how changes in output length affect protein properties at three noise levels: 0.1 (**Figure 3E**), 0.5 and 1.0 (**Figures A.8,A.9**). Raygun introduces short or long indels as needed to achieve the desired length while balancing structural preservation. As expected, larger deviations from the template length led to decreased sequence identity and TM-score. However, this degradation is more gradual compared to noise-induced changes: Raygun largely maintains structural similarity for length changes within ±10% of the template. In specific cases, even larger edits can broadly preserve the structure; e.g., mTOR could be miniaturized by over 500 residues (*>* 20% shrinkage) with a TM-score of approximately 0.7 (**Figure 2E**).

These experiments demonstrate that Raygun’s parameters offer a flexible toolkit for protein design. The noise parameter primarily controls substitution rate, while output length determines the extent of indels. This fine-grained control allows for a spectrum of designs, from subtle tweaks to major structural modifications, with predictable trade-offs between sequence diversity and structural preservation.

#### Raygun compares favorably with existing approaches in balancing sequence diversity and structural plausibility

Raygun’s novel approach to protein design, by introducing substantial indels and mutations in templates, presents a challenge for direct comparisons with existing de novo design methods. To establish a meaningful benchmark, we developed a comparison framework based on sequence masking and regeneration—a task that allows us to compare Raygun with EvoDiff [2], a recent method that can perform de novo design entirely in the sequence space. We mask a portion of the input sequence and task both methods with regenerating the masked regions. This approach allows us to evaluate how well each method balances sequence diversity and structural plausibility when faced with incomplete information. We applied this framework to our set of 20 diverse proteins; for Raygun, we set the noise parameter to 0.5, allowing moderate sequence edits. **Figure 3F** illustrates the results of this comparison.

At 25% masking for both Raygun and EvoDiff, the former improves over the latter across all structural metrics. Since masking impact may not be directly comparable across the two methods, we also evaluated EvoDiff with 10% and 5% masking. At these lower masking rates, EvoDiff’s perplexity, pLDDT, and TM-score become comparable to Raygun’s results at 25% masking. However, EvoDiff’s sequence identity with the original increases significantly at these lower masking rates, indicating less diverse sequences. These results highlight Raygun’s ability to maintain a favorable balance between sequence diversity and structural plausibility. For a given level of structural quality (as measured by pLDDT or TM-score), Raygun generates more diverse sequences. While we do not anticipate this to be the primary way Raygun will be used, this analysis demonstrates Raygun compares favorably with state-of-the-art de novo protein design techniques in its ability to reach new parts of the protein space.

#### Raygun’s fixed-length representations better capture the organization of protein structural shapes

PLMs have demonstrated a remarkable ability to capture fine-grained structural similarities between proteins: at short ranges in embedding space, proteins with highly similar structures cluster together. However, for effective protein design it is crucial that the embedding space also captures global, more high-level relationships. A well-organized representation that captures both fine-grained as well as high-level structural similarities could enable smoother transitions between related structures, facilitating tasks such as protein miniaturization or expansion. Towards this, we investigated whether Raygun’s fixed-length representations improve upon ESM-2 embeddings in capturing these multi-scale structural relationships, particularly for high-level categorizations of protein structure. We leveraged the CATH database, which classifies proteins into hierarchical structural categories, to evaluate the clustering performance of Raygun and ESM-2 representations. Our analysis focused on the three main subsets of the CATH hierarchy: “mainly alpha”, “mainly beta”, “alpha beta”. Within these classes, we focused on the two top levels of the hierarchy: architecture (e.g., “Up-Down Bundle”), and topology (e.g., “Bromodomain-like”).

For architecture and topology levels of CATH, we performed agglomerative clustering using Raygun or ESM-2 average-pooled embeddings. We filtered the dataset for 60% sequence identity and only chose CATH groupings containing at least 50 sequences. **Figure 3C** shows the clustering performance measured by Adjusted Rand Index (ARI) and Normalized Mutual Information (NMI). Both Raygun and ESM-2 representations performed better at distinguishing fine-grained topologies compared to global architectures. However, Raygun outperformed ESM-2 at both levels, with the gap being particularly pronounced at the higher, architecture level.

These results suggest that Raygun’s fixed-length representations not only retain but potentially refine the structural information present in ESM-2 embeddings. The stronger outperformance at the higher level indicates that Raygun captures a better hierarchical organization of protein structure space. This property could be particularly valuable for protein design tasks where preserving global topology is crucial, even while when introducing significant sequence alterations.

#### Raygun demonstrates structural versatility while preserving functional domains

To further assess Raygun’s ability to generate diverse structures at various lengths while maintaining functional integrity, we applied it to preserve PFAM domains across a wide range of structural classes, and input and output protein lengths. We selected PFAM domains corresponding to the four major SCOP classes to ensure structural diversity: *α*-only (7 transmembrane receptor family of GPCR proteins, PF00001), *β*-only (a family of beta-propellers, PF10282), *α/β* (YdjC-like proteins, PF04794), and *α* + *β* (F420 ligase family, PF01996).

For each domain, we chose five templates of diverse lengths and, from each template, generated candidates ranging from 50% to 200% of the median template length in the domain. Our sampling strategy divided this length range into 20 buckets. For each template and length bucket, we initially generated 100 Raygun candidates, resulting in 10,000 (=5 × 20 × 100) candidates per domain. We then applied a filtering step based on *pLL* scores, retaining the top 5 candidates per bucket and template combination, yielding 500 filtered candidates per PFAM domain. To estimate function preservation, we used HMMER to identify candidates that retained their corresponding PFAM domain.

The results demonstrate Raygun’s robustness across structural classes (**Figure 4**). It was able to generate structurally diverse candidates across a broad range of lengths while maintaining functional domains. On average, 48.25% of candidates retained their PFAM domains across all four structural classes. The retention rates ranged from 37% for the most challenging *α*+*β* class to 57% for the *α/β* class, demonstrating Raygun’s robustness even when dealing with complex protein folds. This structural versatility, combined with functional preservation, positions Raygun as a powerful tool for protein engineering tasks that require significant alterations while maintaining the template’s core structural/functional properties.

### 2.4 Generating novel miniaturized fluorescent proteins

We next experimentally tested the ability of Raygun to shrink a class of proteins that have revolutionized cell biology: Fluorescent proteins (FPs) [33], used for over 60 years, have transformed live-cell imaging by enabling the direct visualization of protein localization and dynamics in situ of FP-fusion proteins. Extensive engineering of FPs has produced variants with distinct chromatic properties, monomeric forms, maturation times, and stability, making them indispensable tools for a wide range of applications [9, 30]. However, less attention has been given to reducing their size, a critical consideration for studies involving small proteins, which may be disrupted by their genetic tagging to relatively large FP fusions [36, 29, 46]. Given the central role of FPs in cellular research, and the ongoing efforts to optimize their performance, we tested Raygun’s ability to reduce the lengths of these proteins while preserving their essential function, i.e., fluorescence.

We used Raygun to miniaturize two widely-used fluorescent proteins: eGFP (238 aa), a classic green fluorescent protein, and mCherry (236 aa), a red-emitting fluorescent protein. For each template, Raygun generated four candidates with lengths ranging between 195 and 235 aa. These candidates were selected after filtering 70,000 Raygun-generated samples for each template. The filtering process began with a pseudo log-likelihood (*pLL*) filtering, which eliminated 90% of the samples. Next, we applied the HMM-based tool, *hmmscan*, to remove samples lacking the correct PFAM domain. Finally, we used a custom brightness prediction tool, trained on the GFP brightness dataset [31, 18], to discard samples with low brightness. The detailed sampling procedure is provided in the **Methods A.3.3, A.3.4**. This filtering pipeline resulted in the generation of 8 FPs, XFP01-04 and XFP05-08 being the eGFP and mCherry candidates respectively (**Figure 5**).

**Figure 5:**
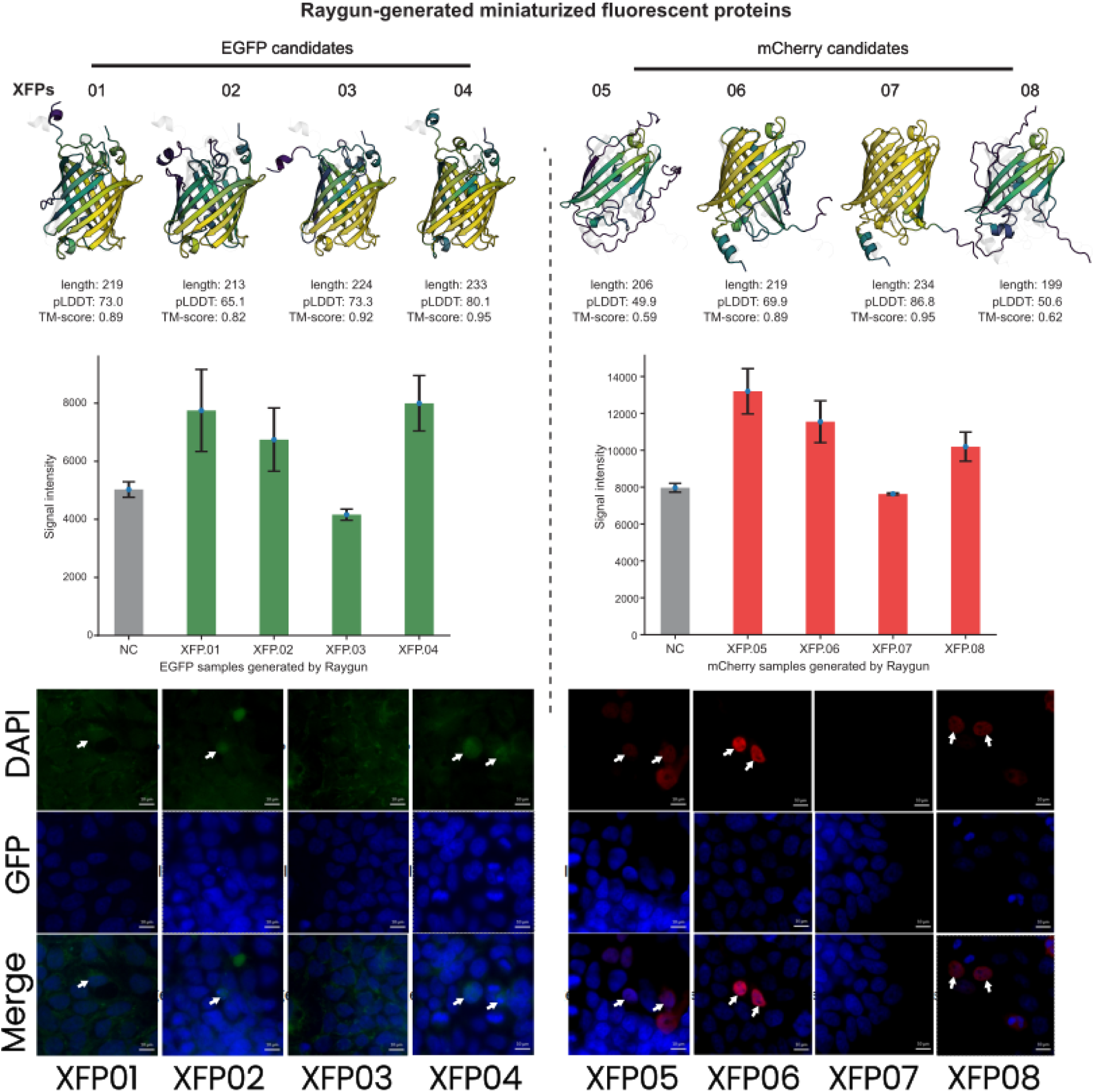
Eight Raygun-generated fluorescent protein (FP) candidates. Four miniaturized candidates each were generated from eGFP and mCherry templates. The AlphaFold-3 predicted structure and metrics (pLDDT and TM-score against the template) for each candidate is shown. The fluorescent images and the corresponding signal intensities for the selected candidates are provided in the succeeding rows. We observed that XFPs 01, 02, 04, 05, 06 and 08 showed significant fluorescence over negative controls, while the XFPs 03 and 07 were confirmed to lack activity.

Our filtering process did not explicitly select candidates based on the presence of the characteristic chromophore sequence, an essential 3 amino acid sequence in the form of *XYG* typically located at positions 65 to 67 in GFP. Recently, Hayes et al. [14] took a scaffolding-based approach to fluorescent protein, including specifying the chromophore. We did not provide any such information to the model: we merely specified the noise and length criteria and let Raygun generate miniaturized FP candidates.

We then proceeded to experimentally test fluorescent properties of XFP01-08. For each candidate, codon-optimized cDNAs were synthesized, cloned into expression vectors and transfected into HEK293 cells. Four days later images were taken and manually inspected for fluorescence over background. Six FPs appeared to display fluorescence over negative control and were selected for further analysis by quantitative image analysis along with two variants that did not appear to have activity. Analysis revealed significant fluorescence over negative control for six of the selected Raygun variants (XFPs 01, 02, 04, 05, 06 and 08), while XFPs 03 and 07 were confirmed to lack activity. Importantly, Raygun was able to generate functional FPs from diverse evolutionary sources. eGFP originates from the jellyfish *Aequorea victoria* while mCherry originates from the coral *Discosoma*. The evolutionary distance between the two groups, Hydrozoa (jellyfish) and Anthozoa (corals), is estimated to be around 600 million years and each FP has markedly different fluorescence properties (such as spectrum). Despite this, Raygun effectively shortened each by up to 25 aa (10.5%) for eGFP and 37 aa (15.6%) for mCherry.

### 2.5 Generating novel miniaturized TurboID proteins

The development of BioID assays—proximity-dependent protein identification using biotin ligases—has revolutionized the field of proteomics, providing a powerful tool for studying protein-protein interactions (PPIs), protein localization and cellular dynamics in ways that were previously infeasible. Originally accomplished using the slow-acting bacterial biotin ligase from Escherichia coli (BirA), engineering efforts have produced improved variants such as BioID2 [17] and TurboID [7]. However, a significant challenge of the BioID approach is that fusion proteins often raise concerns about altering the endogenous properties of the protein of interest. Indeed, around 36% of human proteins are smaller than TurboID, complicating its use as a fusion tag. Reasoning that a shorter sequence may lead to a less intrusive sensor, we explored moderate miniaturizations of the TurboID sequence (1-20% length reduction). Previously, a significantly smaller biotinylating enzyme, UltraID (172 aa), had been synthesized by removing the DNA-binding domain from TurboID. This motivated us to explore if Raygun could autonomously manage large modifications on multi-domain proteins. Accordingly, we applied it to miniaturize TurboID to approximately half its original size and evaluated the resulting design’s sequence, stability and function.

We used Raygun to generate 500,000 TurboID variants, with lengths ranging between 195–235 aa for the moderate-miniaturization objective and 150–180 aa for the extreme-miniaturization objective. Sequence-based *pLL* and HMM screening were then applied, similar to the FP pipeline; the brightness filtering step in the FP generation pipeline was replaced, however, with thermostability-based filtering, enabled by the sequence-based method TemStaPro [28]. Following these sequence-level filters, we screened the remaining candidates through structure-based metrics such as pLDDT and TMscore, finally resulting in 11 candidates: TurboID-{1-10} for the first objective and TurboID-11 (165 aa) for the second. Notably, Raygun autonomously removed the DNA-binding domain to the achieve the 165 aa length, the same domain that was manually removed in UltraID. We emphasize that no domain information was provided to the model and UltraID was not part of its training data. More detail on the candidate generation steps is provided in **Methods A.4**.

Experimental validation of these 11 candidates involved cloning their codon-optimized sequences into expression vectors containing an HA epitope tag for detection. Our experimental screening confirmed that 6 out of the 11 variants, including the TurboID-11 sequence that was shrunk by half, were successfully expressed in HEK cells. To assess their enzymatic activity, we incubated the transfected cells with biotin (overnight, 50 *µ*mol*/L*) and pulled down the biotinylated proteins using streptavidin magnetic beads. Western blotting with an anti-biotin antibody revealed that two variants—TurboID-1 (317 aa, 1% reduction) and TurboID-5 (304 aa, 6% reduction)—exhibited slight enzymatic activity. Therefore, we were successful in our first objective of finding moderately-miniaturized variants with some degree of ligase activity. However, the more difficult objective of miniaturizing the TurboID sequence by half was achieved partially—the expression of TurboID-11 in HEK cells was validated but its ligase active was not significant.

One possible interpretation of the TurboID-11 result is that while Raygun does have the ability of performing large length modifications while retaining poorly understood fine-grained and global interactions required for catalytic efficiency, additional work is needed to a) control the specificity of domain retention, and b) strongly preserve the desired protein functionality. Towards this, a potential solution might involve optimizing the function of the miniaturized design by directed evolution.

### 2.6 Generating EGFR binders using Raygun

The *in silico* design of protein binders is an area of active research with significant potential in drug design and therapeutics. Although Raygun is not specifically optimized for binder design, we sought to evaluate its potential as a general-purpose protein design tool and participated in a protein design competition hosted by AdaptyvBio Inc. Inspired by the CASP competitions, the goal of this competition was to benchmark computational approaches to design protein binders to Epidermal Growth Factor Receptor (EGFR), a target in many cancer therapies. The organizers computationally selected a subset of submissions, synthesized them, and tested them for binding with EGFR. The design constraints were as follows: all candidates had to be 1) single-chain proteins, 2) no longer than 250 amino acids, and 3) at least 10 amino acids different from any published sequence. We participated in the second round of the competition, which was organized following disappointing results from the first round (out of 201 sequences tested, only 5 (2.5%) showed significant binding).

For our design pipeline, we selected EGFR’s endogenous ligand, EGF (53 aa), as the reference template. We then used Raygun, fine-tuned on 5 EGF-like sequences (**Methods A.4**), to generate potential binding candidates. Since EGF’s length was close to Raygun’s minimum-length threshold of 50 amino acids, we opted to modify the template through magnification (instead of miniaturization). We did not impose any explicit constraints to preserve the wildtype EGF binding sites during candidate generation— we wanted to check if Raygun is able to automatically take that into account while generating sequences. After generating 10,000 candidates ranging from 55 to 57 aa, we filtered the sequences to isolate those with the highest predicted EGFR binding potential.

To estimate binding potential, we leveraged ProTrek, a tri-modal PLM trained on sequence, structure, and function data [41]. ProTrek can estimate function from a sequence information alone and we use it to prioritize Raygun-generated candidates. Specifically, we used ProTrek-reported scores for annotations associated with EGFR binding to rank Raygun candidates. Our choice to predict function directly from a PLM—rather than relying on predicted structural metrics like iPTM and iPAE—also distinguished our approach from other methods in the competition.

We submitted 10 EGFR binder candidates, of which 4 entries were selected by the organizers for biological evaluation (referred to as **EGF-Raygun-*`{`*1-4*`}`***; **Methods A.12**). All four candidates were successfully expressed, and two of them (**EGF-Raygun-1** and **EGF-Raygun-2**) demonstrated strong binding to EGFR, with dissociation constants (*K_D_* ) of 0.274 *µ*mol*/L* for EGF-Raygun-1 and 0.561 *µ*mol*/L* for EGF-Raygun-2. Both bound more strongly than the wildtype EGF (*K_D_* = 0.759 *µ*mol*/L*). The AlphaFold-3 predicted complex structures and pharmacokinetic curves for the two binding designs are shown in **Figure 6D**. Analysis of the generated candidates suggests that Raygun may be capturing deeper functional attributes of sequences beyond simple sequence similarity. Among the design approaches that sought to modify or optimize EGF, our approach yielded the best results. Even methods based on the same baseline PLM as Raygun (ESM-2) performed worse. Interestingly, the more successful Raygun candidates had lower sequence identities with the wildtype (EGF-Raygun-1: 70.7%, EGF-Raygun-2: 76.8%), compared to EGF-Raygun-3 (80.4%) and EGF-Raygun-4 (78.6%). Since Raygun effectively identified candidates with higher function and lower sequence identity from a vast combinatorial space of possible modifications, this suggests that Raygun can harness deeper functional characteristics independent of sequence identity while generating candidates.

**Figure 6:**
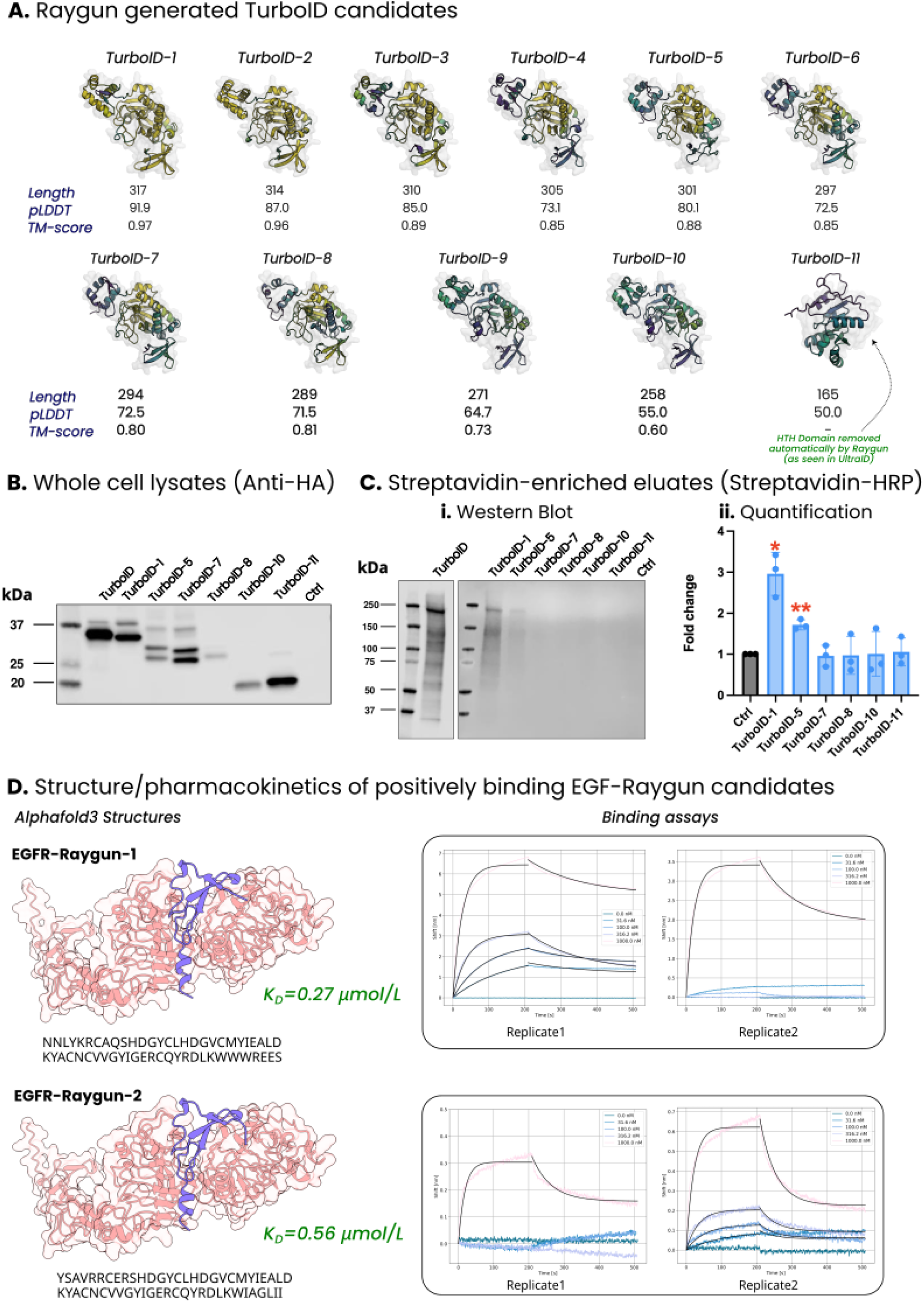
TurboID and EGF binding results: A) Raygun-generated TurboID candidates screened for experimental validation, B) The whole-cell lysate shows expression of 6 out of 11 TurboID candidates in the HEK cells, C) Western blotting results using anti-biotin antibody and corresponding quantification, showing two candidates (TurboID-1 and TurboID-5) with significant biotin ligation activity. D) Pharmacokinetics results and AlphaFold-3 inferred structure of the two EGF-Raygun candidates that showed binding affinities greater than the wildtype EGF.

## 3 Discussion

Raygun introduces a novel approach to template-guided protein design, leveraging probabilistic encoding and language model embeddings to enable extensive modifications while preserving core structural elements. This method allows for both minor tweaks and major alterations, enabling a spectrum of protein variants with high sequence diversity. The capabilities demonstrated by Raygun have far-reaching implications for protein engineering. For instance, we generated functional fluorescent proteins (FPs) with lengths of 199 and 206 amino acids, shorter than 96% of FPs listed in FPbase. Given past research demonstrating the narrowness of FP fitness landscape [31], our Raygun-driven FP optimization—which introduced over 40 indels and substitutions in one case—is particularly significant. We also showed the generalizability of Raygun’s design pipeline through successful miniaturization of the biotin ligase TurboID and optimization of Epidermal Growth Factor (EGF), where we achieved strong binding affinity to EGFR. Notably, the Raygun-designed variants outperformed the native EGF ligand, showing that Raygun can improve upon naturally evolved proteins even in a zero-shot setting.

Raygun’s core innovation lies in representing proteins as probability distributions in a fixed-size embedding space. This approach addresses the challenge of handling variable-length sequences in protein design. By encoding proteins as multivariate normal distributions with fixed parameters, Raygun enables efficient sampling and manipulation of sequences regardless of length. This probabilistic encoding, combined with rich protein language model embeddings, creates a flexible framework for generating diverse protein variants while maintaining core structural properties.

Raygun’s template-based design approach sets it apart from de novo approaches that have been the focus of recent protein design efforts. Template-guided approaches like Raygun offer distinct advantages, particularly in scenarios where modifying existing proteins is preferable due to immunogenicity concerns, compatibility with existing workflows, or the need to minimize in vivo disruption. Moreover, there are certain applications (e.g., miniaturization, or insertion of specific binding, localization, or signaling motifs) where template-guided design is the only feasible approach.

Our experimental validation of FPs suggests that template-guided approaches can, in some cases, enable greater novelty than de novo design. Consider our FP generation approach against a recent de novo GFP design by Hayes et al. Using a new PLM, ESM-3, the latter relied heavily on scaffolding constraints when generating ‘esmGFP’: their approach preserved the length (229) to be the same as the PDB exemplar 1QY3, specified not only the sequence but also the structure of residues critical for chromophore formation (Thr62, Thr65, Tyr66, Gly67, Arg96, Glu222), as well as the structure of residues 58-71, deemed crucial for chromophore energetics. In contrast, we did not impose any such constraints. Notably, most of Raygun’s generated candidates did preserve the chromophore and, given Raygun’s speed, ensuring this for all candidates by explicit filtering would have been straightforward. However, we sought to explore if other chromophores could also work, as this would enable generation of potentially new functionality not available in nature. Therefore, in choosing which candidates to experimentally validate, we included a few with non-canonical chromophores. In particular, the 199-length FP we generated and validated has glycine deleted, its position taken by a serine instead. This demonstrates Raygun’s capacity to explore a broader design space, potentially uncovering novel protein variants that more constrained approaches might miss. In any protein design strategy, there is a balance between staying close to the manifold of feasible proteins while being distinct from the subset of natural proteins. De novo design methods start far from the manifold and try to end at a candidate that resides on it. Raygun, in contrast, starts from the manifold and seeks to stay on it while moving away from the starting point. We speculate that the latter approach may provide a better conceptual balance in reaching novel but feasible proteins.

Broadly, our work suggests that the relationship between sequence and function is more complex than might be currently appreciated, especially for highly engineered proteins like FPs and TurboID. For example, both we and Hayes et al. found that the generated FP candidates are dim and will require directed evolution to reach higher fluorescence. This speaks to the discussion in the field about the kind and amount of experimental training data needed for computational protein design. Towards a low-data approach, Hie et al. have hypothesized [15] that PLM-generated mutations, which have been shown to follow evolutionary rules, are likely to improve fitness in general; they demonstrated this in the case of particular antibodies. This is an appealing hypothesis. If true, it could obviate the need for function-specific experimental data to guide protein optimization. Unfortunately, while this hypothesis may hold true for functions that are evolutionarily rooted, our work with fluorescent proteins suggests that for engineered or non-natural protein functions, the fitness landscape may be more rugged and discontinuous than this hypothesis implies.

Our EGF results provide further insight into this dichotomy. Using a sequence-focused approach without explicit structural optimization, we generated EGF variants with stronger EGFR binding than the native ligand. Notably, the most successful candidates were not those with highest sequence similarity to wildtype EGF. The key sequence modifications occurred at peripheral positions, presumably distant from the binding interface, yet substantially influenced binding affinity. This demonstrates that even for naturally evolved proteins, the relationship between sequence and function can be more complex than structure-based approaches might suggest. Our use of ProTrek for candidate filtering also highlights how function-focused language models can complement Raygun’s generative capabilities, offering an alternative to traditional structure-based virtual screening.

While the evolutionary insights captured by PLMs are very valuable in preserving structure, that may not always be sufficient to guide the optimization of non-natural or highly specialized protein functions. Our TurboID results illustrate this: Raygun autonomously identified and removed the DNA-binding domain whose redundancy (for biotinylation) was only discovered manually through extensive trial and error. While our synthetic miniaturized protein was structurally stable and expressible in cells, its biotinylation activity was modest. This suggests future opportunities in directed domain manipulation— both removal and addition— where computational approaches could be guided by functional constraints. Moreover, it suggests that hybrid approaches that combine the broad exploratory power of methods like Raygun with targeted directed evolution or function-specific experimental assays could be critical for pushing the boundaries of protein engineering.

Our PLM-based approach is very efficient—its single-shot generation takes about 0.3 seconds per iteration on an NVIDIA A100 GPU, about 100-fold faster than the de novo method EvoDiff, which itself is faster than other de novo design methods. This speed, combined with the use of PLM-based evolutionary likelihood filters, enables rapid generation and screening of candidates that deviate significantly from the template while maintaining attractive foldability and structural properties. For instance, in our fluorescent protein experiments, we were able to generate and filter 70,000 samples for each template, applying multiple layers of computational screening before experimental testing. This high-throughput computational approach, coupled with a high experimental hit-rate (5 out of 8 tested FP variants showed significant fluorescence, 2 out of 4 EGF candidates showing enhanced binding affinity), demonstrates Raygun’s potential to accelerate the protein engineering pipelines. Here, we achieved our results using a relatively modest PLM (ESM-2 650M) and a training set of only ∼95,000 sequences. There is significant potential for improvement by incorporating more powerful PLMs, larger training sets, or structure-infused models. The recent advances in PLMs, such as ESM-3 [14] and SaProt [40], suggest that integrating these more sophisticated models could further enhance Raygun’s performance.

Looking forward, Raygun opens exciting possibilities for protein engineering. The novel approach of representing proteins as probability distributions may be useful in other areas of computational biology. Raygun’s ability to efficiently generate diverse, functional protein variants while preserving core structural elements could accelerate drug discovery, particularly for biologic therapeutics, and advance synthetic biology applications such as biosensor design. The framework’s flexibility in handling both substitutions and large-scale indels expands the universe of proteins derivable from a single template, potentially catalyzing new approaches to protein design.

## Declarations

The authors declare no competing interests.

## Acknowledgements

We thank Aditya Parekh, Aditya Pratapa, Huan Liang, Palak Jolly, Sinan Ozbay, Tomoyuki Fujiyama, and Yueshan Liang for assistance in generating structures using the AlphaFold-3 webserver.

## A Methods

### A.1 Model architecture

The “T-Block” layers are arranged before and after the “Reduction” and “Repetition” layers to add model complexity preceding and succeeding the length-transforming operations. In the Raygun encoder, a T-Block output is usually the input to another T-Block and the Reduction layer. The T-Block pairs, that are before and after Reduction, are parameter-shared; we use 10 of them in total. All the reduced T-Block outputs in the encoder are aggregated in the final layer. There, the fixed-length representations are first concatenated together, and then projected to 1280 dimensional space (i.e. the embedding dimension of ESM-2).

Similarly in the Raygun decoder, the T-Blocks are also usually arranged in pairs before and after the Repetition operation, although their parameters are not shared. The target length provided by the user is used to project the fixed-dimensional embeddings back to the variable-length space. All the projected outputs coming from the Reduction layer, after passing through the T-Blocks, are then aggregated in the final projection layer. Similar to encoder, they are first concatenated together before projecting back to 1280 dimensional space. We use 21 T-Blocks in the Raygun decoder: 10 before the Repetition operation and 11 after. The overal Raygun architecture is described in **Figure** A.1.

We now proceed to describe the internal architecture of the Repetition and Reduction layers.

#### A.1.1 Reduction Layer

The Reduction block is a non-parametric layer that transfers variable-length embeddings to a fixed-length space and is vital in the fixed-length sampling process. Given an ESM-2 embedding ***E*** ∈ R*^n×^*^1280^ (1280 being the ESM-2 650M dimension), the Reduction block returns two fixed-length embeddings: mean and standard deviation matrices, each of dimensions R*^K×^*^1280^, where *K* is fixed to 50 in our experiments.

Dividing the variable length embeddings into non-contiguous blocks is tricky if the sequence length *n* is not an exact multiple of the fixed-length dimension *K*. We addressed this by allowing an extra slack of size 1 to the blocks at the beginning and the end of the embedding. For example, suppose the sequence length is 543 and *K*= 50. We chunked the embedding from this sequence by allowing the first 22 and the last 21 blocks to have a size of 11, while fixing the middle 7 blocks to size 10 (22 × 11 + 7 × 10 + 21 × 11 = 543).

We describe the Reduction pseudo-code in **Algorithm 1**

**Figure A.1:**
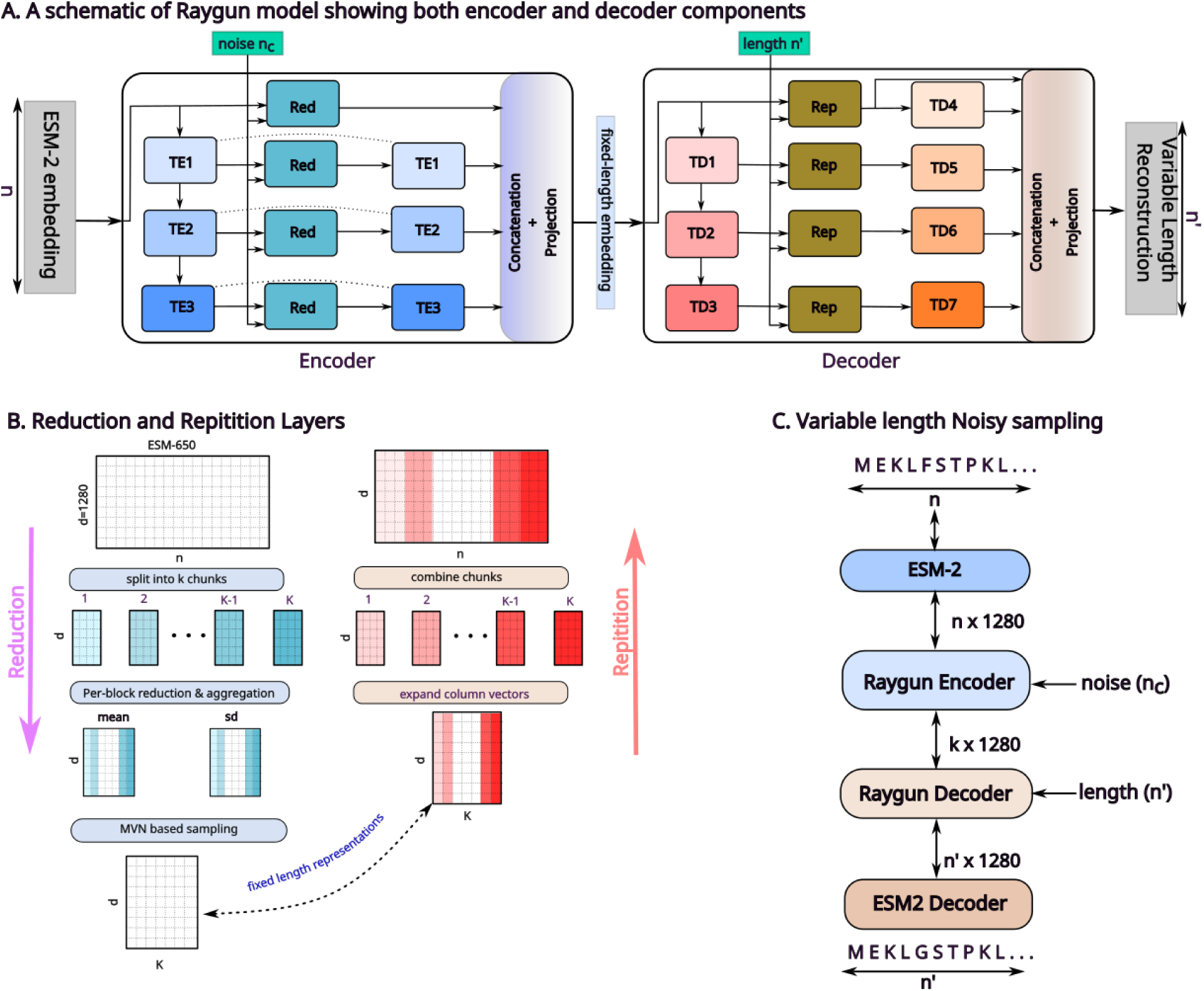
Raygun architecture

#### A.1.2 Repetition Layer

The Repetition layer, which complements the Reduction layer, processes the fixed-length representations from the Raygun encoder and translates them into a variable-length space. This layer performs the length transformation by broadcasting each column vector of the fixed-dimensional representation, ensuring that the resulting combined embeddings have the desired length. Similar to the Reduction layer, the expansion process becomes more complex when the target length *n* is not an exact multiple of *K*. To address this, we allow a portion of the initial and final column vectors to be broadcasted with an additional value of 1. For example, if the target length is 543, the Repetition layer broadcasts the first 22 and the last 21 column vectors into matrices of size R^111280^. The remaining 7 column vectors in the middle are broadcasted into matrices of size R^10^*^×^*^1280^.

The detailed algorithmic description of the Repetition layer is provided in **Algorithm 3**.

##### Algorithm 1 Forward operation of Raygun Reduction Layer

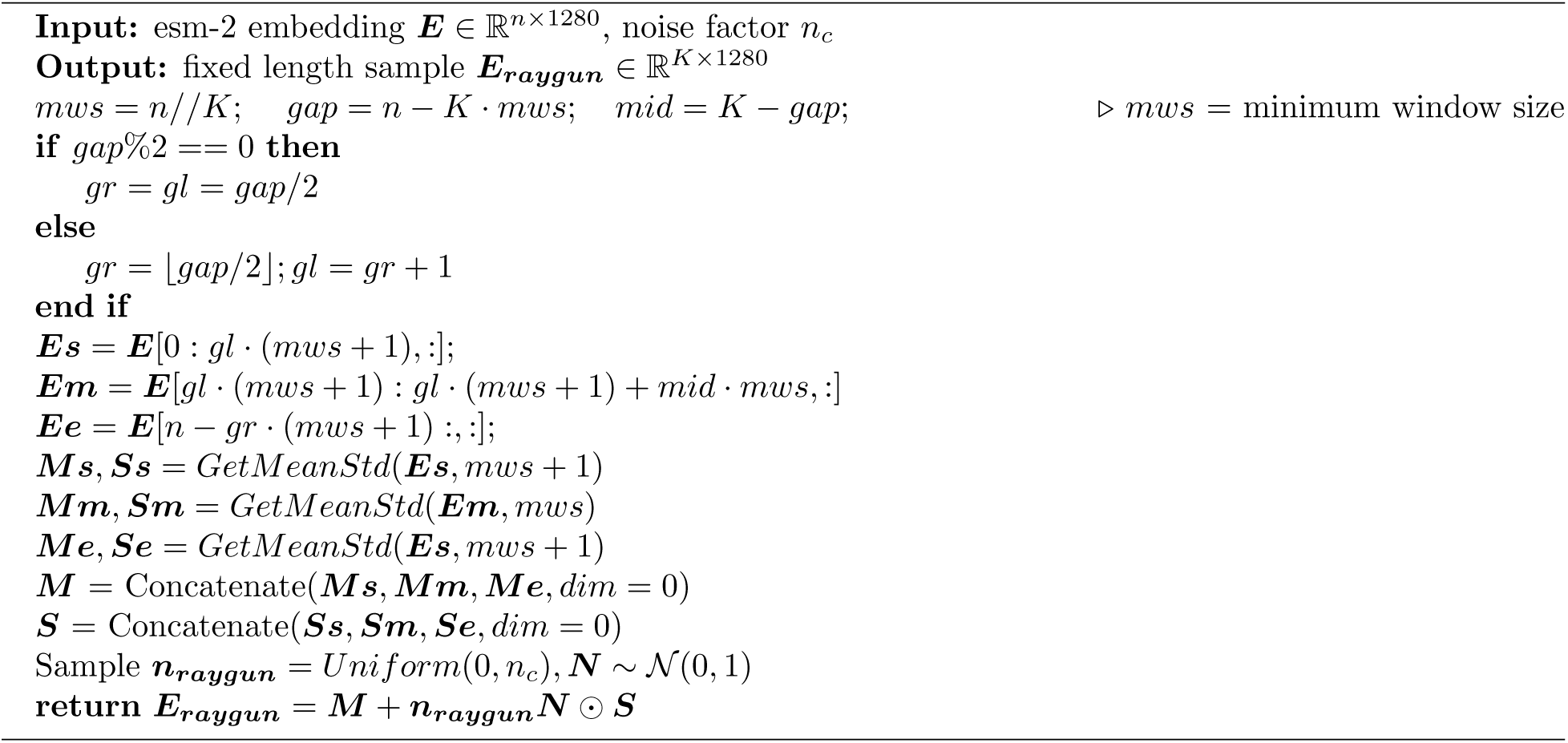

##### Algorithm 2 GetMeanStd() !ht

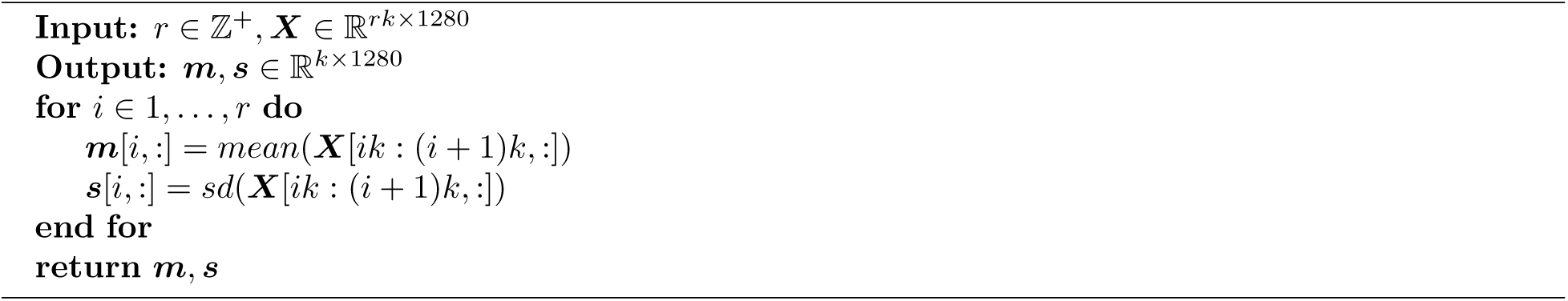

##### Algorithm 3 Forward operation of Raygun Repetition layer

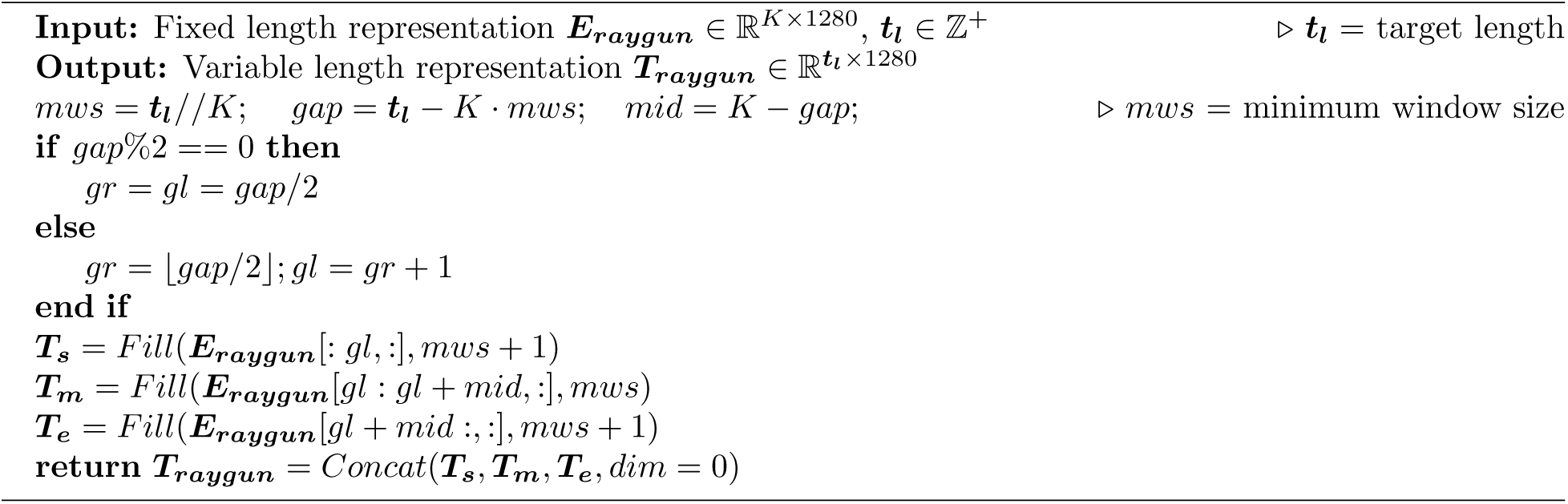

#### A.1.3 T-Block layer

T-Block layers are deep neural network modules trained to optimize embeddings by enhancing latent local and global properties. Each T-Block layer comprises an ESM Transformer for global properties and a 1D-convolution block for local relationships. The internal PyTorch architecture of each T-Block is

##### Algorithm 4 Fill()

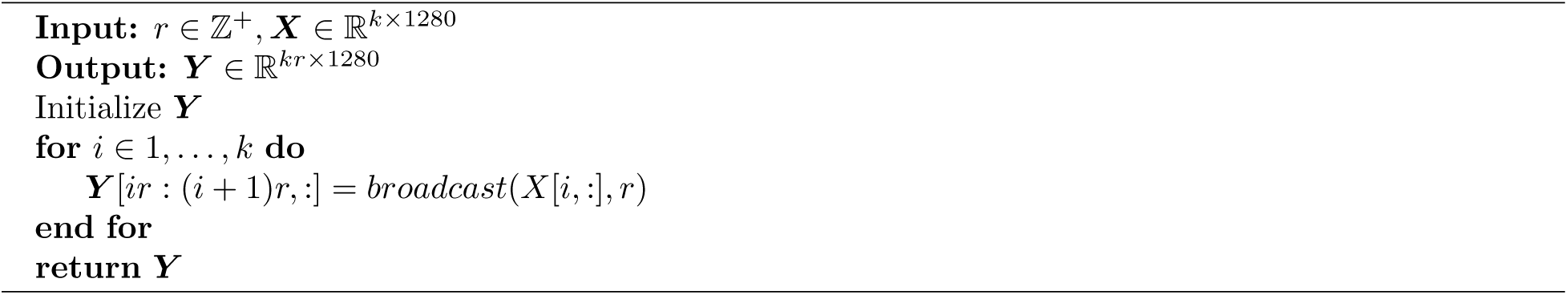

**Figure A.2:**
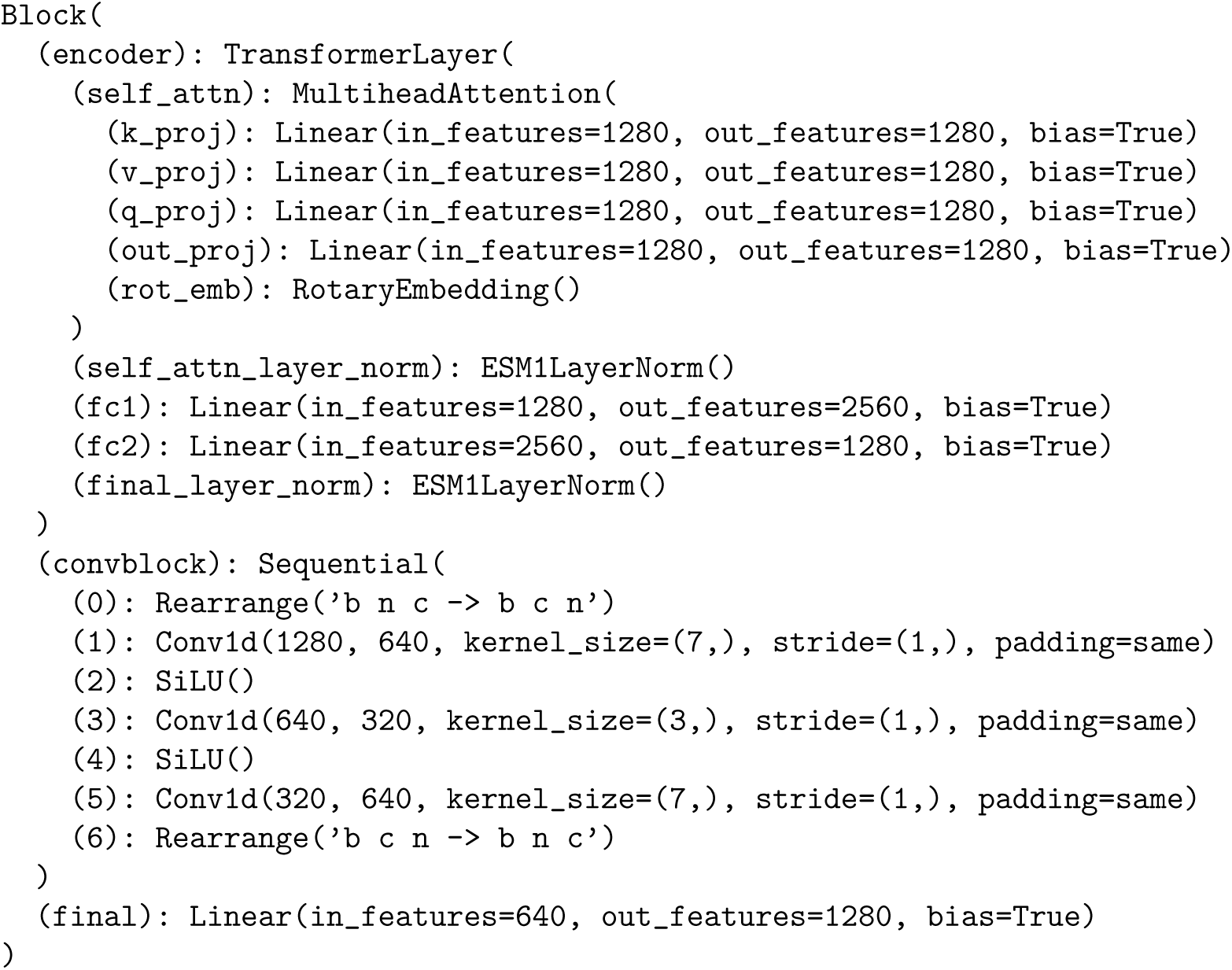
The internal architecture of T-block of the pretrained Raygun model shown in **Figure A.2**.

### A.2 Training Raygun and optimization losses

The Raygun encoder and decoder components are jointly trained to ensure that the decoder can accurately reconstruct the variable-length space from the fixed-length encoding produced by the encoder. To achieve this, we sampled 94,734 proteins from the UniRef50 dataset, carefully maintaining length diversity in the selection. During training, the model takes a protein of length *n*, applies the Raygun encoder to project it into a fixed-length representation where *K* = 50, and then reconstructs the embedding back to its original length *n* using the Raygun decoder. The output from the Raygun decoder is further converted to amino acid logits using a pre-trained ESM-to-tokens decoder. Backpropagation is then performed to minimize the following losses:

1. Cross-Entropy Loss (*L_ce_*): to ensure that the predicted tokens match the input tokens.
2. Reconstruction Loss (*L_rr_* ): to ensure that the reconstructed ESM embedding matches the input.
3. Replicate Loss (*L_rp_*): Let a protein *p* has a length *n* and suppose we generated a new protein *p^′^* of length *n^′^* using Raygun. Then, this loss is designed to ensure that the fixed-length embedding of *p* is close to that of *p^′^*

Let *p* be the input protein of length *n*, *p^′^* the Reconstructed Raygun sequence, *ESM_p_* the ESM-650M embedding of *p*, 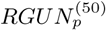 be the fixed length encoding of *p*and 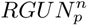 be the reconstruction of using Raygun to length *n*. Let *p^′′^* be another Raygun sequence obtained from *p* of length *n^′^ < n*. Then the total and constituent losses become:

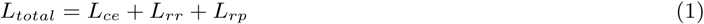

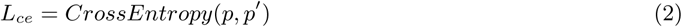

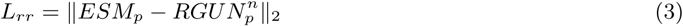

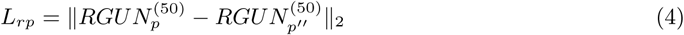

#### A.2.1 BLOSUM score

We used a BLOSUM-based sequence identity score, or simply “BLOSUM score” to evaluate the accuracy of Raygun on the validation dataset. Given the template protein *S* and the predicted Raygun candidate *S^′^*, this score is computed as the ratio:

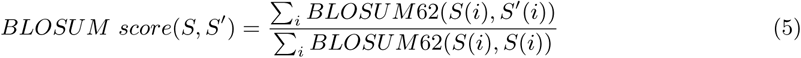

Unlike sequence identity (seq. id), BLOSUM score can range between -1 and 1. Additionally, having a good BLOSUM score is a stronger condition than having a seq. id score of a similar magnitude, as the BLOSUM62 score of two amino acids is usually negative when they do not match.

#### A.2.2 Choice of the reduction parameter *K*

As a preliminary experiment, we aimed to determine the optimal fixed-length reduction parameter *K* for Raygun. To this end, we randomly selected 103,463 sequences from SwissProt, filtered at 50% sequence identity, and split them into 90,524 training and 12,939 validation sequences. We then generated the corresponding BLOSUM scores for *K* = 25 and *K* = 50 after running the models for 3 epochs. Our results showed that *K* = 50 consistently outperformed *K* = 25 in both the training and validation phases

**Figure A.3:**
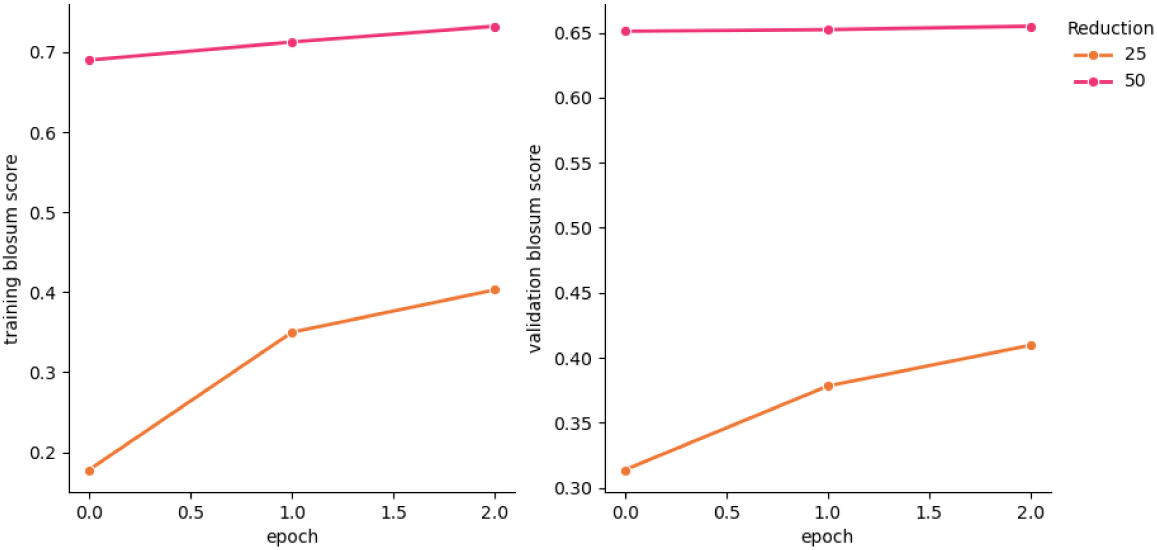
Training and validation BLOSUM scores for K = 25, 50 on the Swissprot dataset (**Figure A.3**). Consequently, we chose *K* = 50 as the default size for the fixed-length representation.

**Figure A.4:**
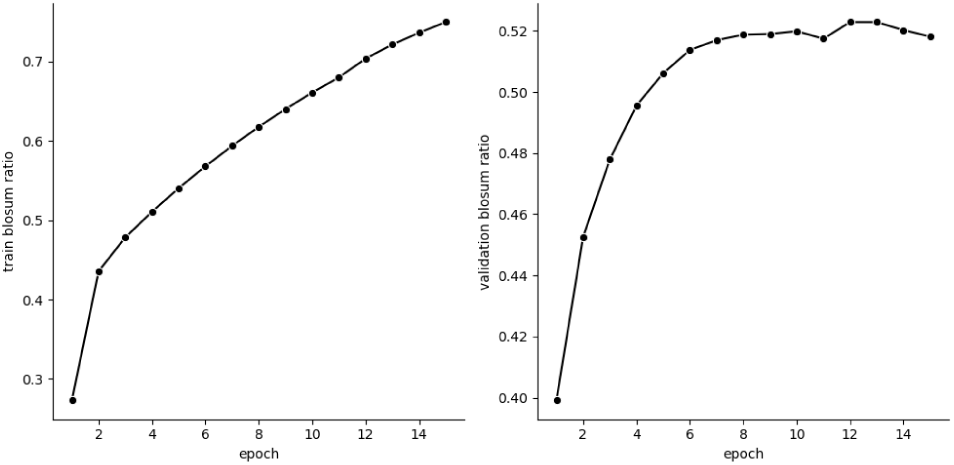
Train and validation BLOSUM scores for epochs1-15

#### A.2.3 Training the model for K = 50

We trained our Raygun model for 15 epochs on 6 A100 GPUs using a Distributed Data Parallel (DDP) approach. Each epoch took approximately 12 hours (8 days in total). Highest validation BLOSUM score of 0.51 was reported on epoch 12. We show the train and test BLOSUM scores in **Figure A.4**.

### A.3 Filtering Raygun samples

The generation speed of Raygun gives us enough flexibility to apply many filtering approaches to improve the quality of the generated candidates. We discuss some of these approaches below

#### A.3.1 pseudo-Log likelihood and repeats penalization

As a first filtering step, we used pseudo-log likelihood [5] to filter the Raygun samples. Given a candidate sequence, *pLL* uses the esm-2 generated logits to compute the degree to which esm-2 identifies the sequence as a viable protein, rather than an arbitrary sequence.

**Figure A.5:**
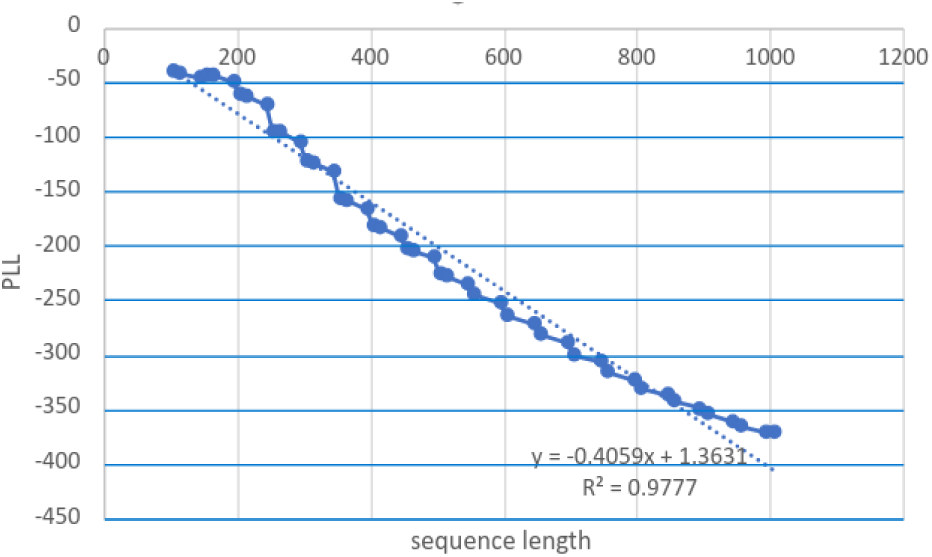
Plot of sequence length vs pLL scores observed for a sample of Uniref50 proteins

One challenge with using *pLL* out of the box is that it is length-dependent and more suited for evaluating substitutions rather than indels. In our generation pipelines, we typically explored multiple output length ranges. Within a particular range, we needed to adjust *pLL* for minor length variations, which we did empirically. We observed an approximately linear relationship between sequence length and the *pLL* score (Figure A.5).

To adjust for length variations, we updated the *pLL* score to make it length-invariant. The updated *pLL_invar_* (*S*), where *S* is the input sequence, becomes

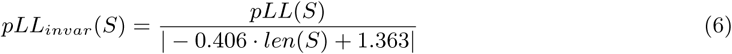

Additionally, in order to discourage sequences with long repeats, we introduced a small sequence-repeat penalty. A similar strategy has been used in other generation methods [2]. We measured the length of sub-sequences with more than 3 repeats *R_rep>_*_3_ with the total sequence length. We then update *pLL_invar_* again using *R_rep>_*_3_ in the following way:

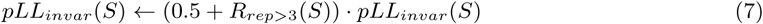

This updated metric, which is always negative, is then used to find candidate sequences with higher fitness (i.e. having *pLL_invar_* values closer to 0).

#### A.3.2 Using HMMER for evaluating PFAM domains

In PFAM-based experiments (Section 2.3 of the main paper), we used HMMER [11] as an evaluation tool to determine the percentage of generated Raygun samples that retain the original PFAM domain of the template. After *pLL* filtering, the Raygun candidates for each PFAM domain were processed through HMMER. The ratio of candidates that retained the domain after length modifications was then used to evaluate Raygun’s ability to preserve domain information in its generated samples.

**Figure A.6:**
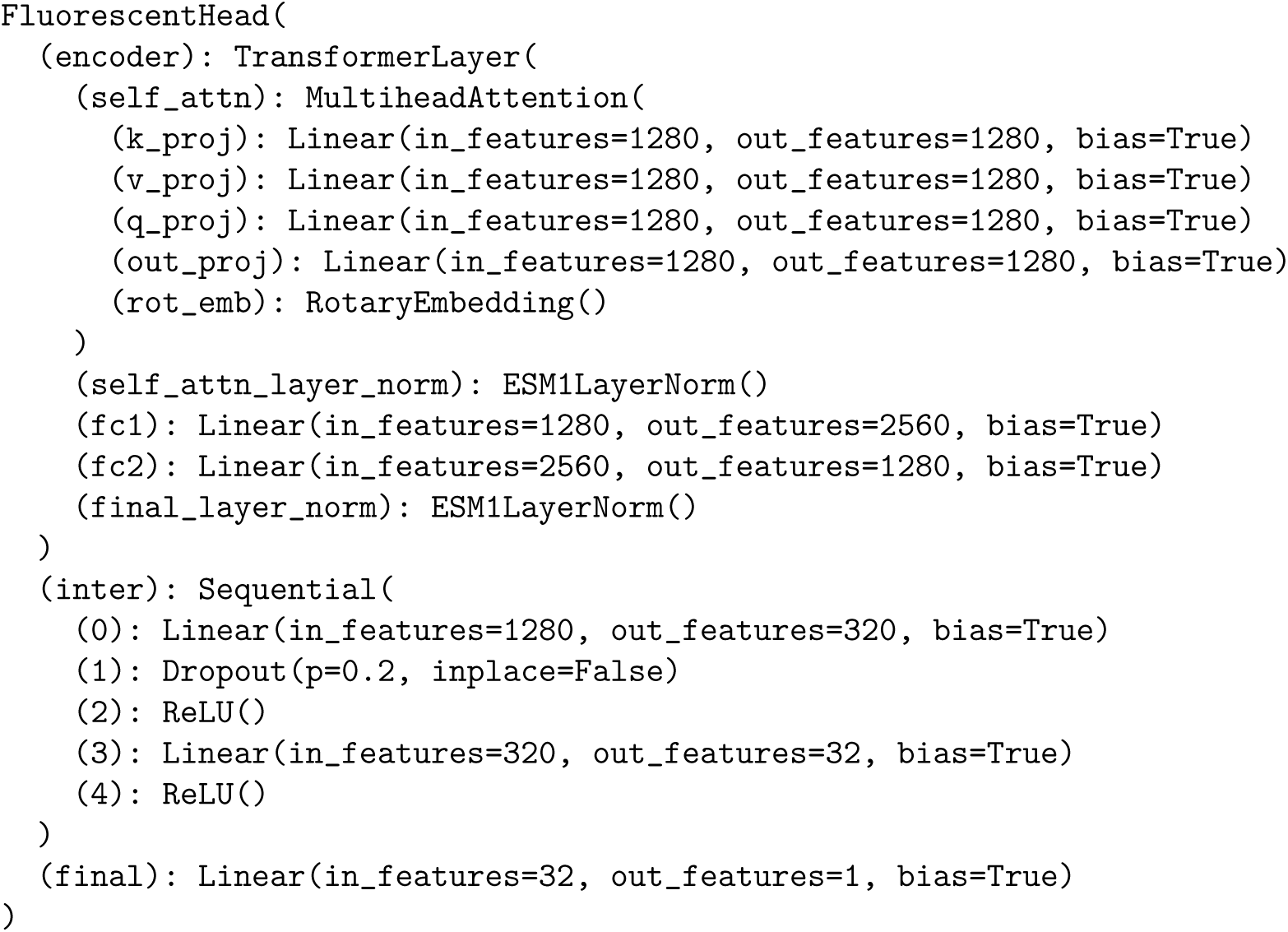
Internal architecture of the Fluorescent head

#### A.3.3 Using HMMER as a filtering tool for fluorescent protein generation

In the fluorescent protein experiments, we used HMMER as an additional filtering tool to discard Raygun-generated candidates lacking the appropriate PFAM domain. For new FP generation, we specifically checked if the candidates belonged to the PFAM domain “PF01353”. The PFAM score threshold was set to 20 during filtering.

#### A.3.4 Using known GFP brightness information for additional FP filtering

In addition to *pLL* and HMMER-based filtering, we also used existing brightness data obtained from a deep mutational scan (DMS) assay on avGFP [31] to train a deep model and select candidates predicted to be the brightest. The DMS dataset contained brightness results for 54,025 sequences. We constructed a simple fluorescent head on top of ESM-2 (with its weights frozen) and trained it to predict the brightness values of input DMS variants of avGFP. The internal architecture of the fluorescent head is shown in Figure A.6. We trained this model for 50 epochs.

When deploying this trained brightness model, we aimed to ensure that it would not negatively impact sequence diversity by selecting a narrow range of sequence diversity. Therefore, we first clustered the candidates filtered by *pLL* and HMMER into 15 disjoint clusters. We then applied the brightness model to each of the 15 clusters and obtained the 15 brightest predicted sequences for both eGFP and mCherry. The clustering was done as follows:

1. For the remaining candidates, perform Multiple Sequence Alignment (MSA) using MAFFT [16].
2. Compute pairwise BLOSUM scores between all candidate pairs aligned in the MSA. Use these pairwise similarity scores to construct an affinity matrix.
3. Finally, use spectral clustering to group the candidates into 15 non-overlapping clusters.

In the final stage, we generated AlphaFold3 structures for these 30 candidates in total and used the obtained PDBs to compute TM-score and pLDDT. We used this structural information to manually select 4 candidates for each FP, ensuring that candidates across diverse lengths were included. The amino acid sequences of these 8 candidates are provided in **Figure A.7**.

#### A.3.5 Advantages of Recycling

Although Raygun can be used to perform one-shot generation, we hypothesized that the diversity of the generated candidates would be greater if we applied a one-step recycling procedure: taking Raygun output and passing it back again to the encoder, in order to get a recycled product. Recycling has been shown to be a powerful tool to enhance the reach of a model without adding additional model complexity, and has been widely used in methods like AlphaFold2 and ESMFold. To test this hypothesis, we took 5 proteins from the Rhodopsin family (PF00001): *ADA1D HUMAN*, *CCR1 HUMAN*, *FSHR BOVIN*, *LPAR6 CHICK* and *NTR HUMAN*, as templates, fine-tuned Raygun on these 5 proteins and used them to generate 38 candidates per template, spread over the length ranges 90 to 1010. We set the Raygun noise-factor to 0.5, and used *pLL* as a filtering step. We then used HMMER to estimate the percentage of these candidates that retained the “PF00001” domain.

We performed this experiment for the “no-cycle” and “1-step recycle” settings, and found that the number of candidates with the PFAM domain retained after recycling was significantly higher at 68*/*190 than the candidates generated without doing any recycling (50*/*190). Therefore, the default invocation of Raygun for protein generation includes 1-step recycling, which can be also be disabled by the user.

**Figure A.7:**
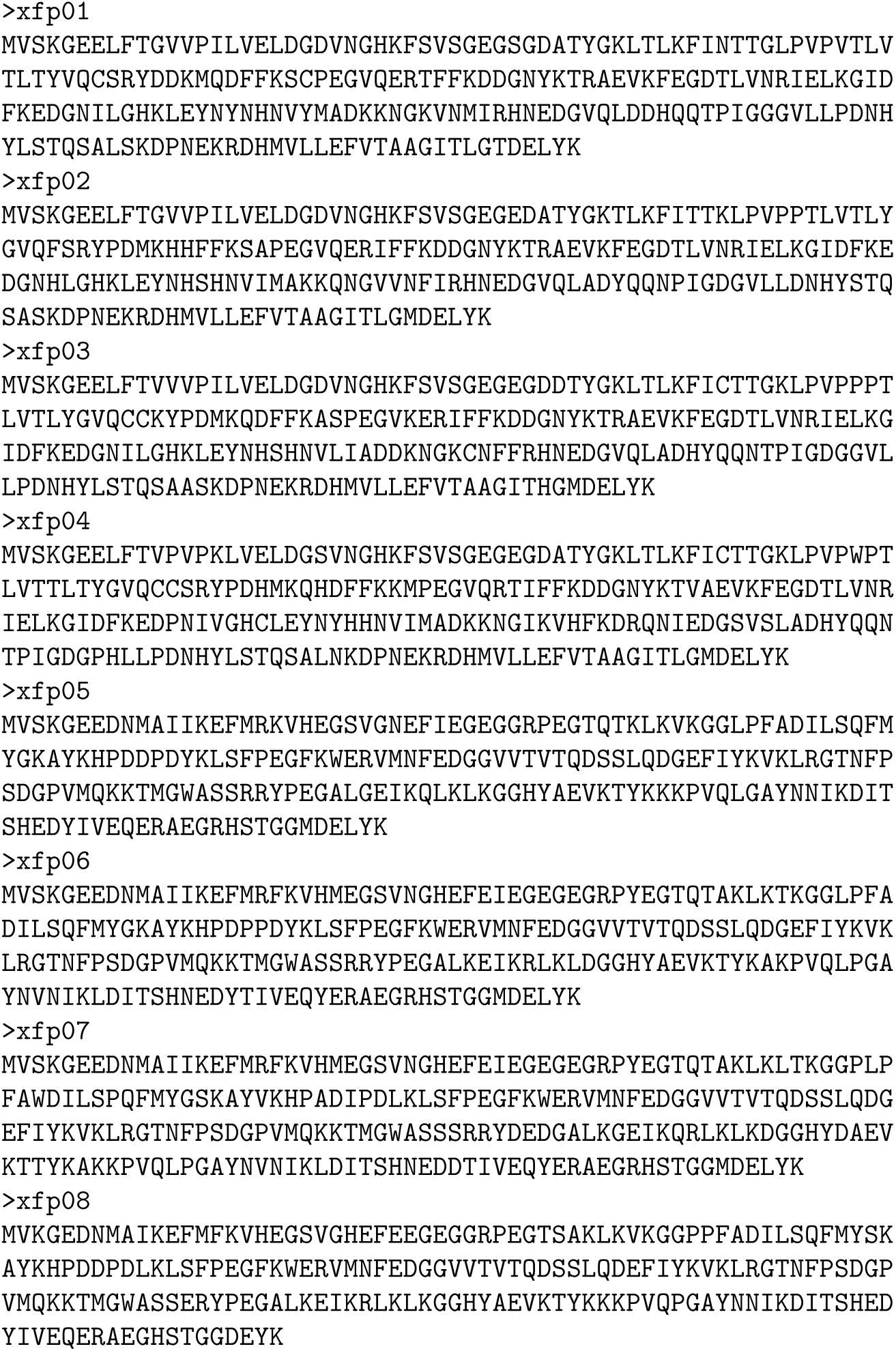
Amino acid sequences of the 8 Raygun-generated XFP candidates

**Figure A.8:**
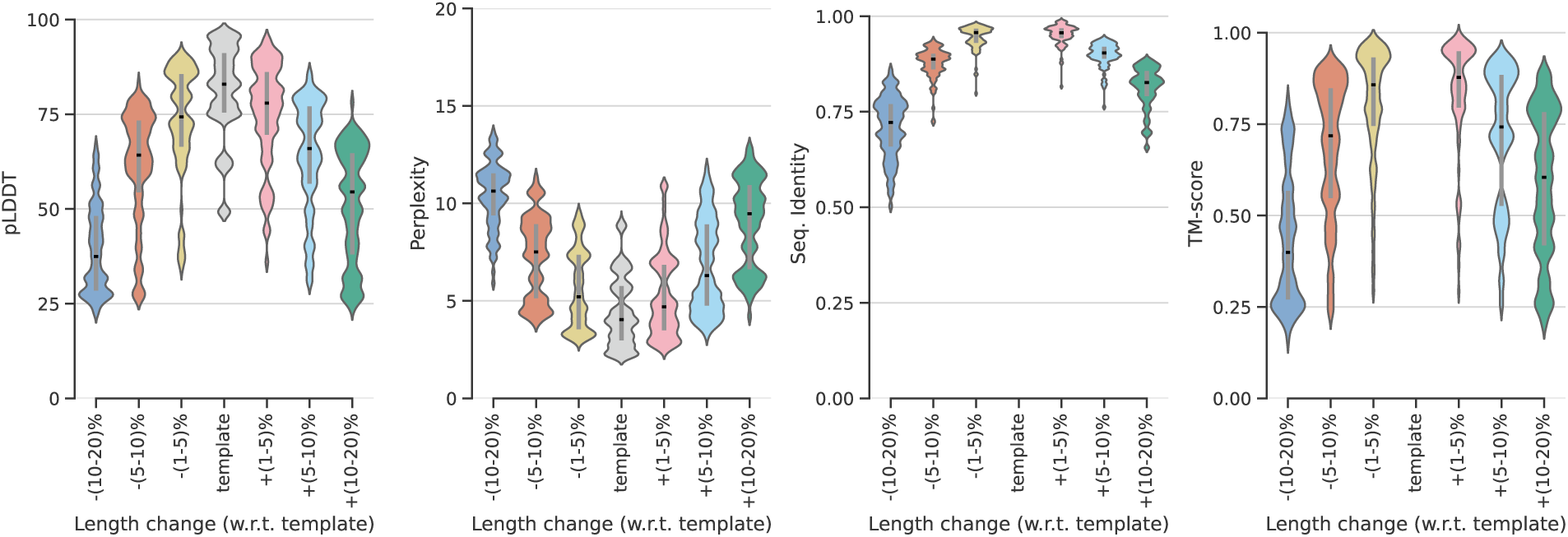
pLDDT, perplexity, seq. identity and TM-score results, after changing candidates lengths by different margins, for noise-factor 0.5

**Figure A.9:**
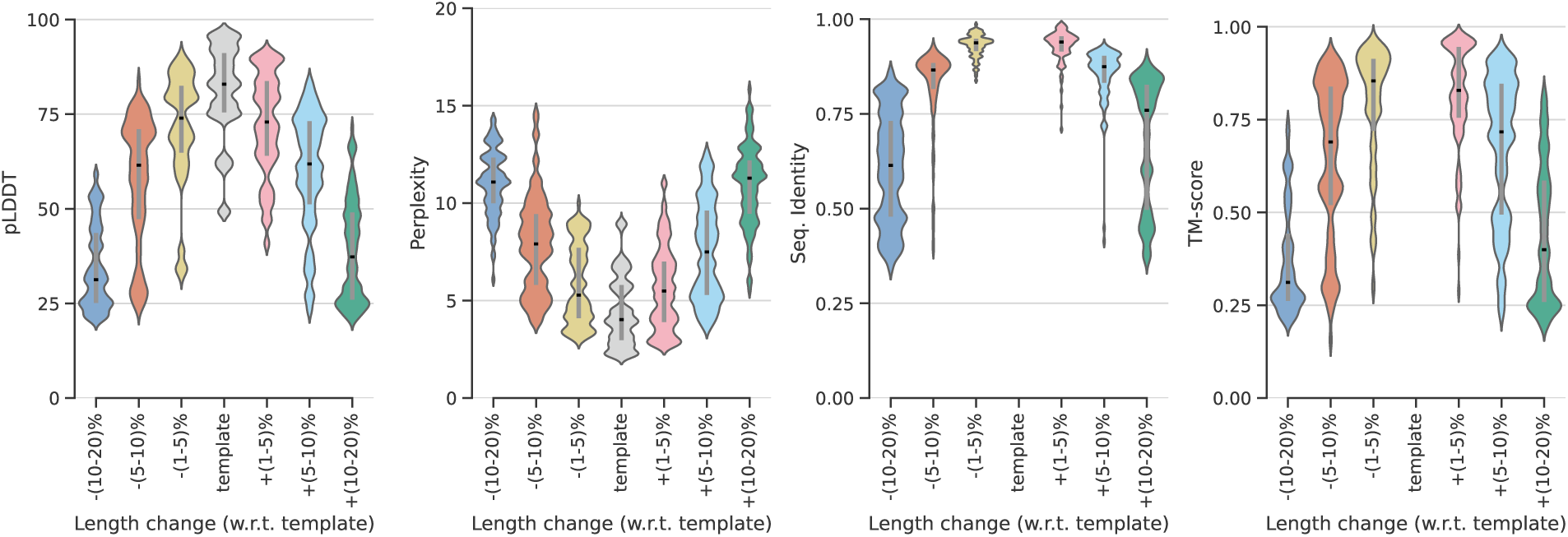
pLDDT, perplexity, seq. identity and TM-score results, after changing candidates lengths by different margins, for noise-factor 1.0

### A.4 TurboID candidate selection process

Our initial steps involved fine-tuning model to perfectly reconstruct the TurboID, UltraID and the original BirA sequences. For the first goal of finding larger-than-UltraID variants, the model was used to generate 500,000 TurboID candidates between the ranges of 240 to 320. The subsequent filtering steps proceeded in a similar way as our FP pipeline, with the length-adjusted *pLL* being the first filter, selecting only 1 out of 5 candidates. The brightness filtering step in FP generation, however, was replaced with a ‘thermal stability’ module, which used the sequence-based model “TemStaPro” model [28] to predict the thermal stability of the candidate sequences, thereby selecting only half the candiates obtained after *pLL* filtering. Subsequently, we applied hmmscan to remove the candidates that did not retain two PFAM domains: ‘PF03099’ and ‘PF02237’, associated with biotin ligation. In the final sequence-based filtering steps, we sorted the generated candidates based on how well they retained the known TurboID binding sites, after pairwise alignment with the original TurboID sequence. After the application of MAFFT-based clustering (similar to **Methods A.3.4**), the overall sequence-based filtering process resulted in 50 possible candidates. These candidates were then subjected to structure based filtering to find candidates with high pLDDT and TMscore values, finally resulting in 10 candidates (TurboID-{1-10}) for final experimental validation (**Methods A.10**). Additionally, towards the second objective of exploring large length modifications, we used the pipeline above to generate and test one candidate of length 165 (referred to as “TurboID-11”).

**Figure A.10:**
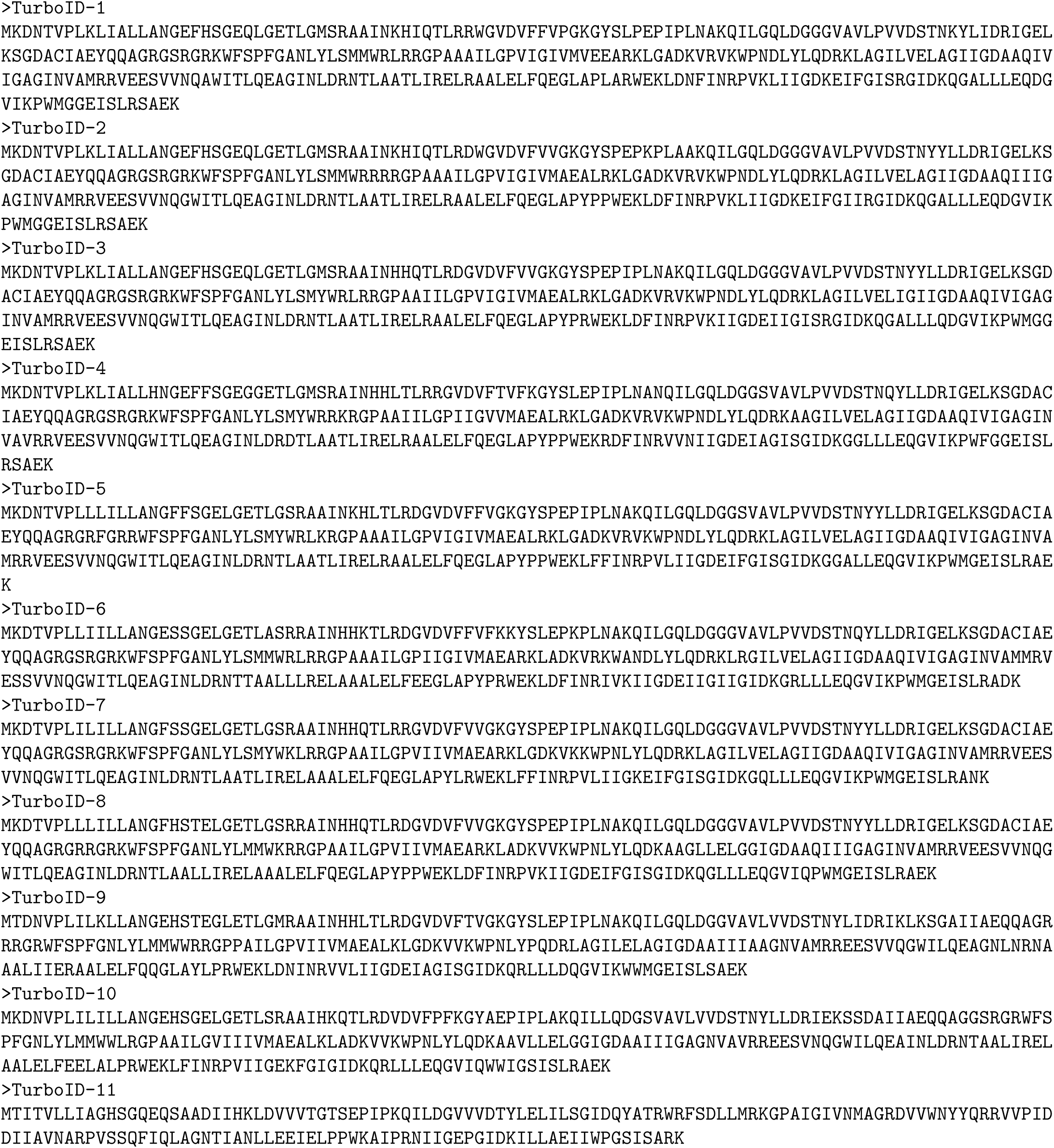
Amino acid sequences of the 11 Raygun-generated TurboID candidates

**Figure A.11:**
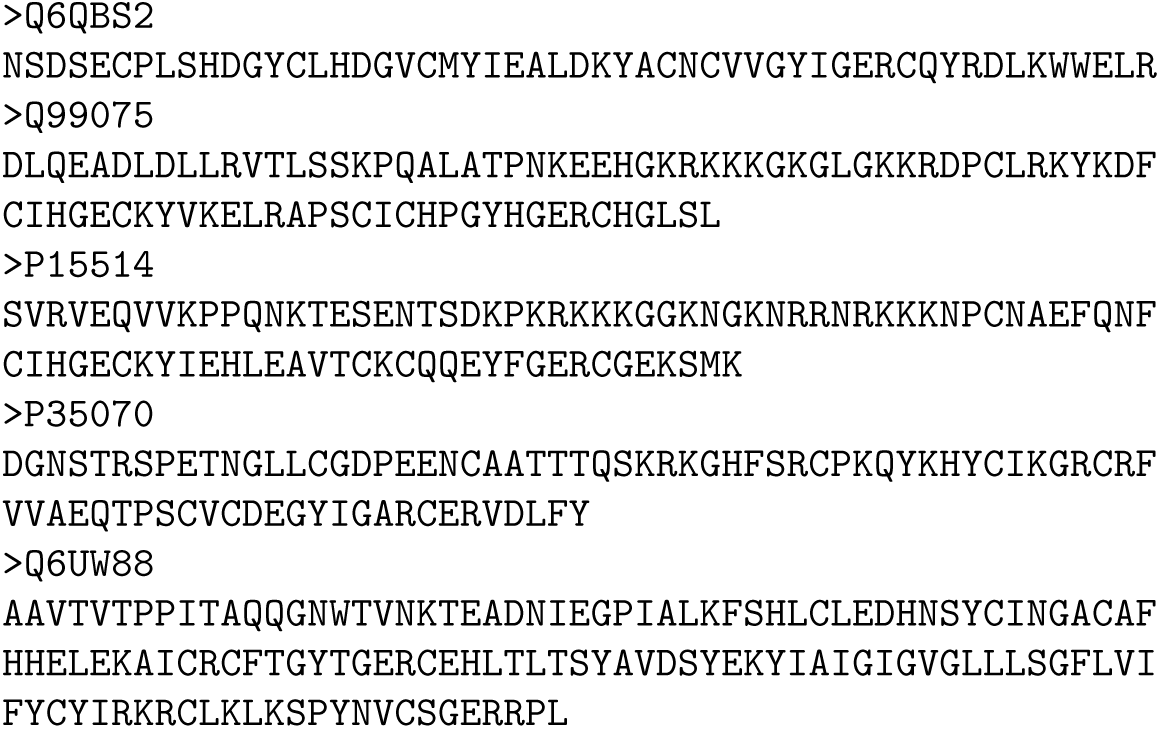
5 EGF-like sequences used for fine-tuning the model for EGF generation

**Figure A.12:**
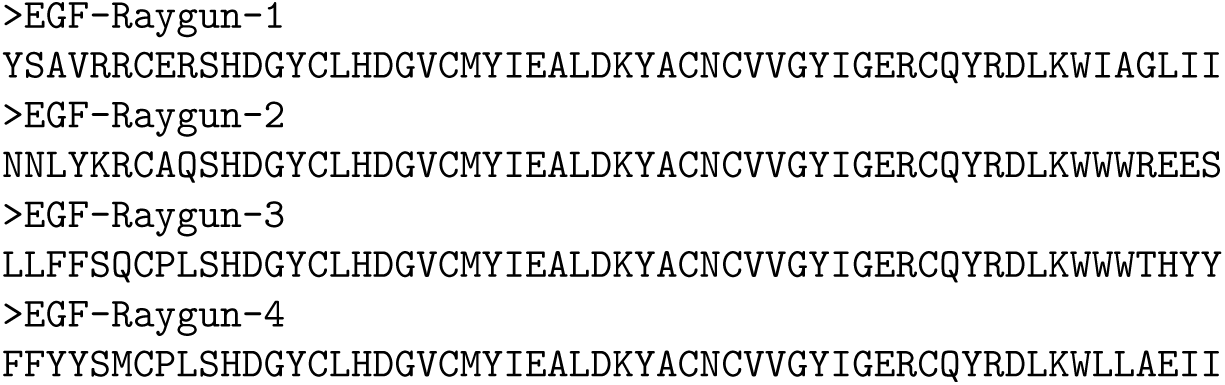
4 Raygun-generated EGF candidates submitted for validation in the Adaptyvbio competition

